# Learning the Unseen: Data-Augmented Deep Learning for PTM Discovery with Prosit-PTM

**DOI:** 10.1101/2025.11.07.687302

**Authors:** Wassim Gabriel, Daniel P. Zolg, Victor Giurcoiu, Omar Shouman, Polina Prokofeva, Florian Seefried, Florian P. Bayer, Ludwig Lautenbacher, Armin Soleymaniniya, Karsten Schnatbaum, Johannes Zerweck, Tobias Knaute, Bernard Delanghe, Andreas Huhmer, Holger Wenschuh, Ulf Reimer, Guillaume Médard, Bernhard Kuster, Mathias Wilhelm

## Abstract

Post-translational modifications (PTMs) are critical regulators of protein function, yet confidently identifying and localizing PTM sites across proteomes remains a challenging task. Integrating peptide property predictions into spectrum interpretation improves identification performance, but training data enabling zero-shot prediction across diverse PTMs are scarce. Here, we present a major expansion of the ProteomeTools dataset, comprising over 977,000 synthetic peptides, covering 22 PTM–residue combinations. Furthermore we developed Prosit-PTM, a model with chemically-informed encoding and amino acid substitution-based augmentation trained with our novel ground-truth dataset, that achieves accurate zero-shot predictions. Applied to modified peptides, Prosit-PTM enhances PTM-site localization in phosphoproteomics, increases identification of multiply modified peptides in histones, and enables data-driven rescoring for unseen modifications such as HLA peptides. Furthermore, the learned embeddings of amino acids and modifications capture physicochemical relationships underlying PTM-driven HLA presentation. Prosit-PTM is integrated into multiple open-source tools enabling PTM-aware rescoring, site localization, spectral library generation, and beyond.

## Introduction

Protein post-translational modifications are key regulators of protein function and cellular processes^1^, with their dysregulation linked to many diseases^2–4^. Tandem mass spectrometry (MS/MS) is the leading approach for large-scale post transitional modification (PTM) identification^5^. Continuous technological advances and improved enrichment methods have substantially increased the ability to detect and characterize PTMs, resulting in the identification of approximately 260,000 modification sites in the human proteome to date^6^. However, identifying PTMs in peptides presents a significant challenge for traditional database search engines^7,8^, primarily due to the combinatorial explosion of the search space from including PTMs and the lower spectral quality resulting from their substoichiometric abundance. Recent deep learning models including Prosit^9–11^, MS2PIP^12,13^, and pDeep^14–16^, have advanced peptide retention time and MS/MS spectral predictions but remain constrained to unmodified peptides or a limited set of supported modifications. While subsequent models, such as DeepLC^17^ and AlphaPeptDeep^18^, can predict properties of modified peptides not included in the training data (unseen PTMs) without the need of re-training (zero-shot learning), they show substantial bias toward seen PTMs with much poorer performance on unseen PTMs. At this time, it is unknown whether this is due to model architecture choice, selected encoding strategy, or limited availability of training data. Because of limited availability of accurate zero-shot prediction models, rescoring platforms such as Oktoberfest^19^, Inferys^20^ or MS²Rescore^21^ can either not be applied or are less effective without (manual) transfer learning of models to PTM-rich datasets. As a result, identification of diverse PTMs remains a challenge, necessitating zero-shot PTM-aware prediction models in proteomics. This study introduces a substantial extension to the synthetic ProteomeTools dataset, covering essentially all frequent biological modifications and addressing the lack of high-quality training data. This dataset was used to developed Prosit-PTM, a deep learning model designed to overcome generalization limitations in PTM characterization. The model employs improved chemically informed PTM encoding which, in combination with a novel amino acid substitution-based data augmentation strategy, allows scaling the training data to virtually unlimited size and thereby enabling the coverage of a much larger chemical space. With this, Prosit-PTM achieves precise zero-shot predictions for unseen PTMs. Crucially, it maintains high accuracy when applied to different complex datasets, addressing a critical gap in PTM proteomic analysis.

## Results

### Extension of ProteomeTools to synthetic post-translational modified peptides

ProteomeTools is an established initiative that has generated and systematically analyzed comprehensive libraries of synthetic peptides based on the human proteome using multimodal liquid chromatography–tandem mass spectrometry (LC-MS/MS). Initially, it started with tryptic peptides^22^, then expanded to include non-tryptic peptides^10^ and later incorporated tandem mass tag (TMT)-labeled peptides^11^. Although we have previously demonstrated the value of synthetic modified peptides in the 21PTMs^23^ and citrullinated peptides^24^ datasets, these efforts have been limited by the relatively small size of available datasets.

Here, we significantly scaled up these efforts with the introduction of ProteomeTools-PTMs. The dataset comprises eight distinct PTM, encompassing 15 unique residue-PTM combinations (Figure 1a). Additionally, peptides with four post-translational modifications (phosphorylation, acetylation, ubiquitination, and methylation) were selected for reanalysis after TMT-labeling, and a large portion of the unmodified peptides were reanalysed after dimethyl labeling. The resulting peptides were organized into 33 packages, each mainly containing a specific type of PTM or specialized groups to address particular challenges, such as phospho site localization on permutations or different alkylation modifications on Cysteine or multiply modified peptides (Methods). The final dataset comprised 977,000 synthesized peptides, consisting of 300,000 TMT-labeled modified peptides, 377,000 unlabeled modified peptides, 250,000 dimethyl-labeled peptides, and 50,000 unmodified counterparts (Supplementary Figure S1a). Multimodal LC-MS/MS data were acquired as described previously^22^, using 5 different fragmentation methods in DDA mode (Methods). Subsequently, MaxQuant was used for identification, confidently assigning ∼50,000,000 peptide-spectrum matches (PSMs), corresponding to ∼700,000 labeled and unlabeled peptides (Figure 1b).

**Figure 1.**
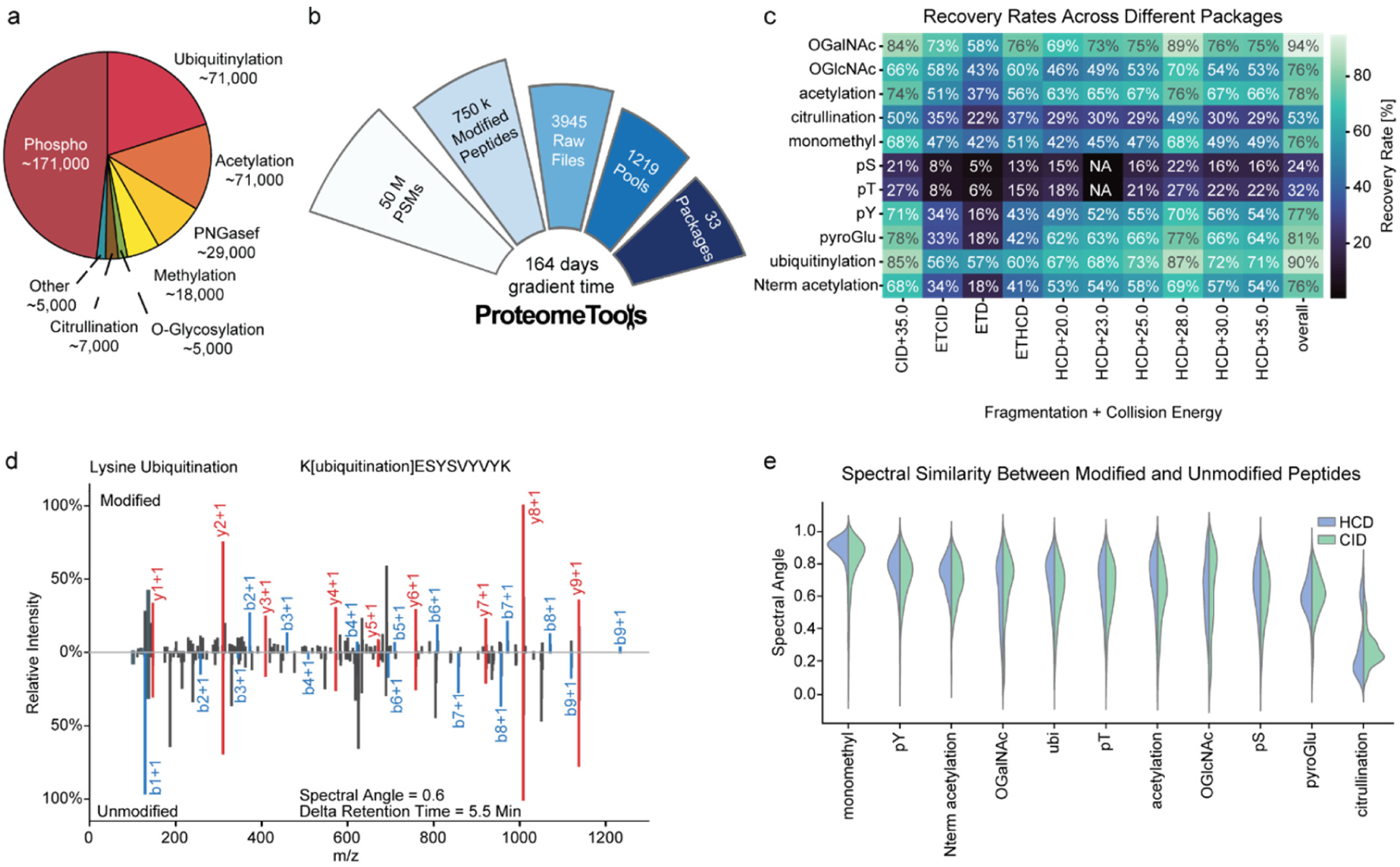
Recovery and Spectral Characteristics of Modified Synthetic Peptides in ProteomeTools-PTMs. **a** Distribution of post-translational modifications (PTMs) in ProteomeTools-PTMs. The pie chart shows the distribution, with counts of synthesized modified peptides for each distinct PTM. **b** The dataset consists of 36 million PSMs derived from ∼750,000 modified peptides extracted from 3945 raw files. Peptides were divided into 1219 sets of at most 1000 peptides each across 33 differently themed packages (e.g. acetylation, ubiquitination etc.). The analysis required a total gradient time of 164 days. **c** The heatmap displays relative recovery rates (color coding) for different residue-PTM combinations (y-axis) analyzed under various mass spectrometric fragmentation conditions (x-axis). **d** Mirror spectrum comparison of modified and unmodified peptide experimental spectra. The spectrum displays the MS/MS fragmentation patterns of ubiquitinated peptide K[ubiquitination]ESYSVYVYK versus its unmodified counterpart KESYSVYVYK. The blue and red peaks represent annotated b-ions and y-ions, respectively. **e** Violin plots of spectral angle (SA) values between modified and unmodified peptides across different modifications and fragmentation methods (HCD blue, CID green) which highlight the variability in the experimentally acquired spectra, with some modifications producing highly similar spectra and others leading to pronounced spectral divergence.

Due to the variation in fragmentation patterns of modified and non-modified peptides^23^, we employed 10 distinct acquisition parameters for each pool (Methods). To identify the most effective method for each modification, we analyzed the synthesis recovery rates across all modifications. The results revealed that O-GalNAc-modified peptides achieved the highest overall recovery rate at 94%. In contrast, the recovery rates for phosphoserine (pS) and phosphothreonine (pT) were significantly lower, at 24% and 32%, respectively (Figure 1c), which is largely due to lower synthesis yields for phosphorylated peptides. Generally, normalized collision energy (NCE) of 28 for HCD fragmentation resulted in maximal identifications on our instrument compared to lower and higher NCEs.

The effects of PTMs on fragment ion intensities (FII) and indexed retention time (iRT) have been previously examined on a small scale within a subset of Proteometools^23^, analyzing only 200 unique peptides, both with and without PTMs. To validate these findings, we expanded the analysis to over 100,000 unique peptides. Notably, fragment ions that do not carry the modification also showed small variations in intensity, but the majority comes from ions that carry the modification (Figure 1d). To determine whether this effect is limited to a single PSM or is more widespread across the dataset, we analyzed spectral similarity based on annotated ions without accounting for *m/z* differences; there remains a significant difference in the FII, with an average spectral angle (SA) of 0.6 (Figure 1e). A similar pattern is observed for iRT where the average delta iRT between modified peptides and their unmodified counterpart is 6.6 (Supplementary Figure S1c). This suggests that FII and iRT predictions for unmodified peptides are not reliable for evaluating modified peptides.

In summary, ProteomeTools-PTM is the largest publicly available systematic (near) ground-truth dataset for PTMs available to date. It provides a rich and systematically characterized resource of modified synthetic peptides and, in combination with previously generated data, lays the groundwork for developing more sophisticated computational tools capable of accurately predicting and identifying PTMs.

### Enabling zero-shot prediction of unseen PTMs via synthetic peptides and encoding strategies

Using the complete ProteomeTools resource, we extended our generic artificial neural network architecture (Methods), termed Prosit (Latin for ‘of benefit’), to enable high-accuracy predictions for iRTs and FIIs of modified peptides. For this purpose, multiple ProteomeTools datasets^9–11,22^ were used for training, including unmodified peptides, TMT-labeled peptides, dimethyl-labeled peptides and the modified peptides added here (Methods). The previously published 21PTMs dataset<u>(Zolg et al. 2018)</u> was reserved as an additional holdout dataset, containing novel modifications to assess model generalizability. For the model architecture, we decided to follow the Prosit architecture closely but added an additional encoder for PTMs (Figure 2a, Methods).

**Figure 2.**
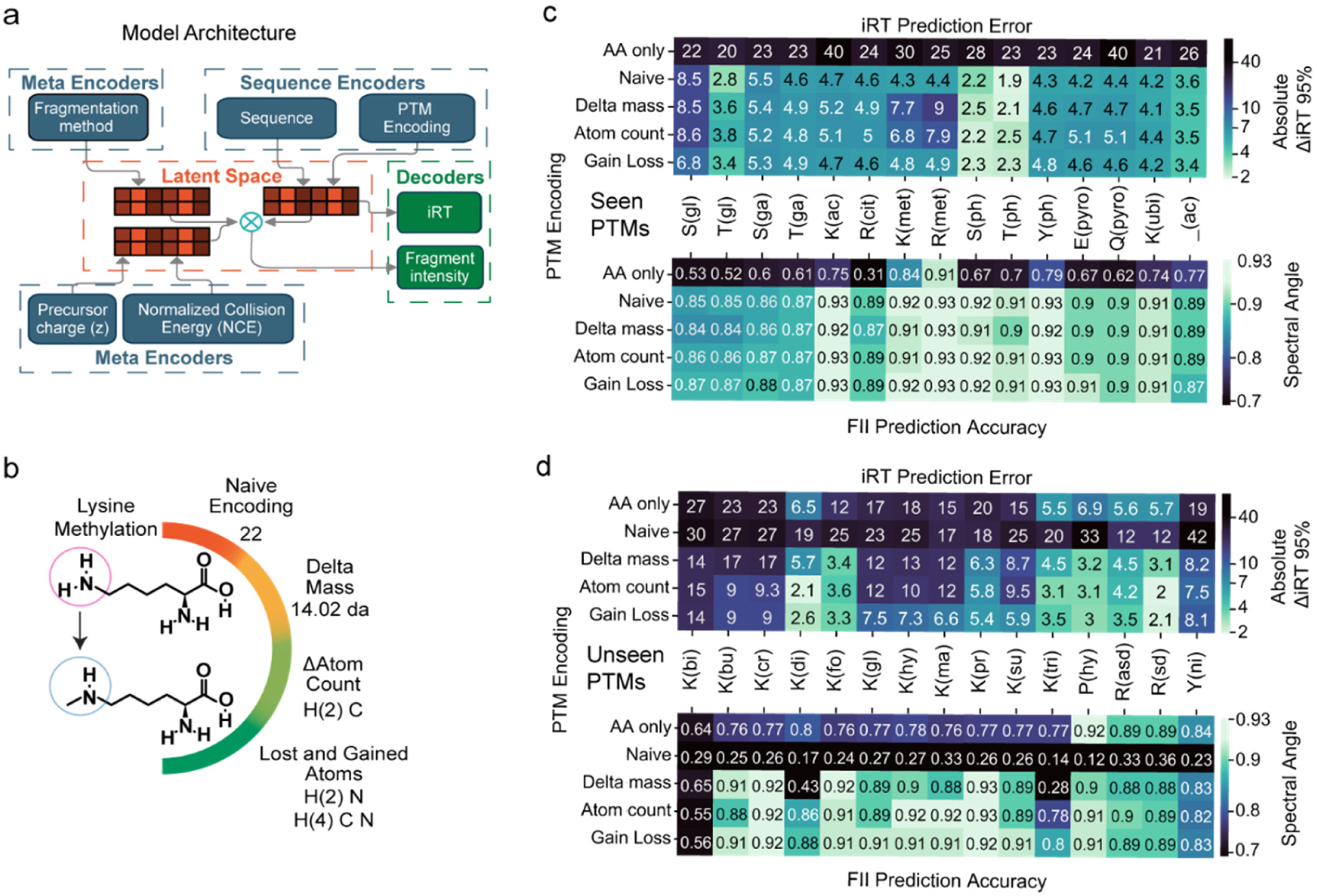
Comparative Analysis of PTM Encoding Strategies in Training and Model Performance. **a** Schematic representation of a multi-encoder neural network architecture is presented to predict peptide properties from mass spectrometry data. This model integrates information through three primary components: Sequence encoders for the amino acid sequence and PTM features, meta encoders for experimental conditions (fragmentation method, precursor charge, and normalized collision energy), and decoders for predictions. The indexed retention time (iRT) decoder utilizes information from the sequence encoders only, while the fragment ion intensity (FII) decoder uses the sequence encoders and meta data encoders. **b** Exemplary illustration for lysine methylation of the different PTM encoding methodologies used , arranged in order of increasing detail. The naive encoding strategy (orange) uses a static number (also referred to as tokens) for each modification. In contrast, the delta-mass approach (yellow) accounts for the mass change caused by the modification in relation to the unmodified residue. Additionally, the delta atom count (light green) describes differences for C, N, P, S, O, and H atoms. The final method explicitly specifies the gained and lost atoms separately (dark green). **c** Comparison of PTM encoding techniques in Prosit-PTM for predicting modified peptides with seen PTMs (present in the training set) on both iRT and FII. The top panel shows a heatmap of the 95th percentile delta iRT errors for iRT predictions. The bottom panel shows a heatmap of the spectral angle for FII predictions. Lower values (lighter green) indicate better predictions. **d** Comparison of PTM encoding techniques in Prosit-PTM for predicting modified peptides with unseen PTMs (not present in the training set) on both iRT and FII. The top panel shows a heatmap of the 95th percentile delta iRT errors for iRT predictions. The bottom panel shows a heatmap of the Spectral angle for FII predictions. Lower values (lighter green) indicate better predictions.

There are multiple conceivable strategies to encode PTMs. Thus, we trained four different models using increasing levels of PTM encoding detail (Figure 2b) and compared their performance for PTMs present in our training data (seen PTMs) and PTMs not present in our training set (unseen PTMs). First, the naive encoding is to assign a unique numerical value (token) to each of the 22 amino acid-modification combinations. Although such a model cannot generalize to PTMs not present in the training set, it is expected to achieve maximal prediction performance on the training data. Second, encoding a PTM by its delta mass (Δmass) enhances generalization by incorporating the mass change associated with modifications (e.g., 14.02 Da increase for methylation). Alternatively, PTMs can be described by the net number of atoms (Δatom) gained and lost (e.g., +H(2)C for methylation). Last, the most descriptive approach tested here encodes separately the atoms gained and lost (gain-loss, e.g., +H(4)CN and -H(2)N for methylation).

For seen PTMs, all encoding strategies showed high prediction accuracy. Using naive encoding, 95% of the predicted versus experimental iRT differences (ΔiRT95) were within 4.3 iRT units, and predicted to experimental FIIs demonstrated strong agreement with an average SA of ∼0.9 (Figure 2c). Increasing the level of encoding detail (Δmass, Δatom count, or gain– loss) did not improve predictions for modifications seen in training, indicating that naive encoding is already sufficient for these cases. For unseen PTMs, performance varied depending on the encoding strategy. The Δmass encoding improved predictions for some modifications, such as K[Malonylation], but failed to distinguish between modifications with similar mass shifts but different FIIs (Supplementary Figure S4c), as seen with trimethylation (Δmass = 42.0469 Da) with a median SA of 0.28, since it has a similar mass to acetylation (Δmass = 42.0157 Da), which was in the training data. The Δatom count encoding offered an improvement over the Δmass approach. Gain–loss encoding produced the most reliable predictions, with average ΔiRT95 of 6.1 and average SA of 0.87 across unseen PTMs (Figure 2d). Still, accuracy declined for modifications with unique atomic compositions not included in the training data, such as biotinylation (ΔiRT95 = 14 and SA = 0.56). Because iRT prediction accuracy improved significantly with gain–loss encoding, and FII prediction showed a marginal improvement, we selected this encoding for further model development and evaluation.

While the generalization capabilities of iRT and FII predictions can be improved by the choice of the PTM encoding strategy, there appears to be further need for data that encompasses a broader range of variability in chemical space. The results suggest that diversity in training data is essential for accurately predicting unseen PTMs in a zero-shot manner.

### Accurate Generalization to Novel Modifications via Amino Acid Substitution-Driven Data Augmentation

While ProteomeTools-PTM is the largest resource of synthetic modified peptides and covers most biologically relevant PTMs, many of the 1,568 described modifications in Unimod^25^ still lack high quality training data. This limited coverage likely restricts the model’s generalizability. In other scientific fields, data augmentation is a common approach to overcoming such limitations^26^, but such approaches are not readily available in proteomics. However, as nearly all amino acids (AAs) can be represented by substitutions from other AAs, we hypothesized that using these substitutions during training can expand the chemical space presented to the PTM encoder and thus enhance generalization.

To test this, we downloaded all available 342 AA substitutions (AASs) from Unimod. We aimed to systematically replace AAs in our training data with one of the available AAS, augmenting the available experimental data. For this purpose, we employed two approaches (Figure 3a; Methods). Firstly, we replaced unmodified AAs with corresponding modified counterparts (the AAS). For example, we replace the unmodified aspartic acid (D) without a modification by asparagine (N) with the PTM Asn->Asp (shown as N[+O-HN] to the model; Supplementary Figure S5a). Secondly, we addressed the substitution of already modified AAs. To illustrate this process, consider a methionine (M) that was oxidized (shown as M[+H(3)CO-H(3)C] to the model). In this case, for example, we first substitute the methionine by an alanine and add the modification Ala->Met (shown as A[+H(5)C(2)S-H] to the model). After this substitution, we further incorporate the existing modification by adding the oxidation state, resulting in ‘doubly’ modified residue (A[Ala->Met][oxidation] shown as A[+H(5)C(2)OS-H] to the model; Methods). In both approaches, the experimental spectrum and observed retention time can be used twice for training, annotated once with the original modified peptide sequence and once with the (hypothetical) AAS modified peptide sequence. While the resulting molecules are chemically identical, those are not shown directly to the model but rather the separated base sequence and modification pattern, requiring the model to learn what the resulting molecule is from the individual components. By utilizing these two approaches, we expanded the chemical space from the previously 22 amino acid-modification combinations to 364, a more than 15-fold increase. Because different AAS can exhibit varying effects on peptide properties, when comparing the iRT or FII of the respective peptide with the base sequence (Figure 3b; Supplementary Figure S5b), it is essential to carefully sample the AASs to ensure to capture AAS that would lead to minor or drastic changes across all peptide properties (Methods).

**Figure 3.**
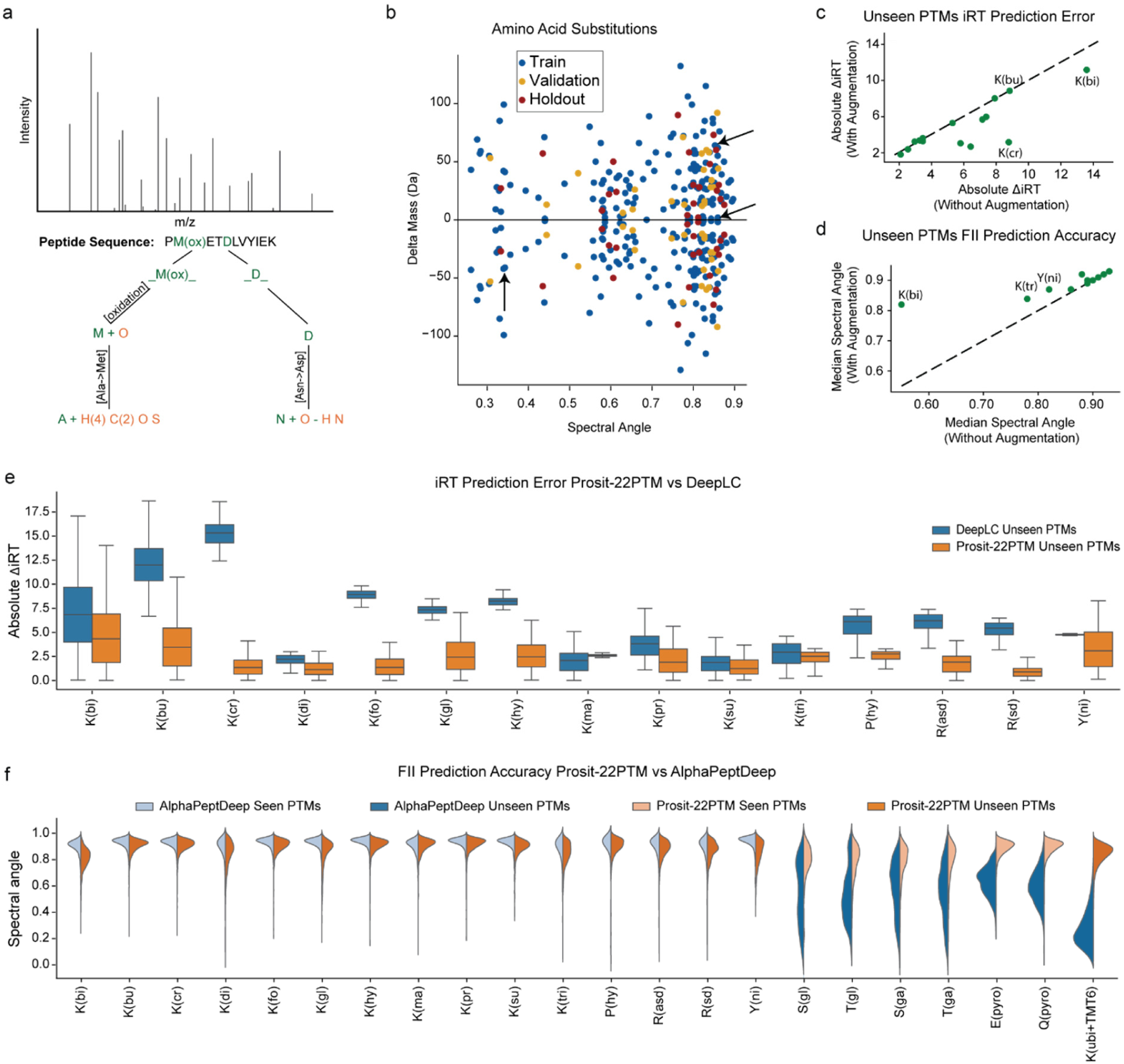
Data Augmentation with Amino Acid Substitutions Enhances PTM Prediction Performance. **a** Schematic illustration of the peptide sequence augmentation strategy. The top panel shows a representative MS/MS spectrum for the peptide sequence PM(ox)ETDLVYIEK, where M(ox) signifies oxidized methionine. The middle panel displays the peptide sequence with the modified residue highlighted. The graph outlines potential augmentation pathways: modified or unmodified amino acids (e.g., Met, Asp) can be replaced with alternative amino acids, along with a modification. The PTM encoding shown to the model is written in orange color. **b** Scatter plot shows individual amino acid substitutions, with points colored by dataset partition: green for training set, blue for validation set, and red for holdout set. The x-axis indicates the spectral angle, reflecting the similarity between MS2 spectra, while the y-axis shows the delta mass (Da) associated with each substitution. **c** Scatterplot shows the iRT delta 95% values for all unseen PTMs across Gain Loss models with and without augmentation. The three PTMs with the lowest accuracy on the models without augmentation are highlighted. Lower values indicate better predictions. **d** Scatterplot shows the spectral angle (SA) for all unseen PTMs across Gain Loss models with and without augmentation. The three PTMs with the lowest accuracy on the models without augmentation are highlighted. Higher values indicate better predictions. **e** Boxplot shows the distribution of absolute iRT differences for predictions of unseen PTMs by both DeepLC (blue) and Prosit-22PTM (orange). **f** Distribution of spectral angles between predicted and experimental MS2 spectra for peptides with PTMs, comparing AlphaPeptDeep and Prosit-22PTM. Light blue and orange indicate spectral angles for peptides with seen PTMs. Dark blue and orange represent unseen PTMs for AlphaPeptDeep and Prosit-22PTM, respectively.

Training data augmentation led to an average 30% increase in accuracy for iRT predictions and a 12% increase for FII predictions on challenging, unseen PTMs (e.g., biotinylation). The previously chosen gain-loss model reduced ΔiRT95 predictions by up to 55% for Crotonylation compared to models trained without augmentation (Figure 3c). These improvements are likely due to the model’s expanded chemical space and balanced representation of modified and unmodified PSMs (Supplementary Figure S4c). For the FII prediction model, using the augmented training set achieved an average SA of 0.89 for unseen PTMs, with a 49% improvement in biotinylation due to the inclusion of sulfur-containing modifications from AAS involving Methionine (Figure 3d). To confirm that including modifications with sulfur (as a result of an AAS) causes the improvement on biotinylation, we trained models on two different datasets: one with only AAS that contain sulfur atoms (Prosit-WA-Su) and another with all other AAS but the ones with sulfur atoms (Prosit-WA-No-Su). The Prosit-WA-Su clearly outperformed Prosit-WA-No-Su on biotinylation, with a 51% improvement, while showing similar or worse performance on other modifications (Supplementary Figure S5c). The gain-loss model trained with augmentation, now referred to as Prosit-22PTM (22 amino acid– modification combinations), demonstrated superior performance and was utilized for subsequent analyses.

Benchmark evaluations against state-of-the-art methods revealed that Prosit-22PTM outperformed DeepLC in iRT prediction for PTMs not included in either model’s training set, achieving over 30% higher prediction accuracy (Figure 3e). For FII prediction, Prosit-22PTM model demonstrated high accuracy on unseen PTMs, achieving a SA of ∼0.9 in several instances, a level of accuracy comparable to that of AlphaPeptDeep on seen PTMs (Figure 3f). In contrast, AlphaPeptDeep’s performance significantly declined (SA ∼0.5) when evaluated on unseen PTMs, while Prosit-22PTM maintained strong performance (SA ∼ 0.89). Only one PTM, TMT6-labeled ubiquitinylated lysines, was unseen by both models, where Prosit-22PTM demonstrated substantially better performance (SA ∼ 0.85) over AlphaPeptDeep (SA ∼ 0.25). Using AAS to augment training data, particularly for PTMs, significantly improved model accuracy in predicting properties of unseen modifications. This method addresses the lack of high-quality experimental data on PTMs and supports zero-shot learning. It not only surpasses existing state-of-the-art tools but also paves the way for expanding the chemical space of any peptide property predictor.

### Enhanced Sensitivity and Precision in Phosphopeptide Site Localization

Accurate localization of modifications, particularly for phosphorylation, remains a critical challenge, especially for peptides containing adjacent serine, threonine, or tyrosine residues^27^. With Prosit-22PTM’s accurate iRT and FII predictions at hand, we hypothesized that this limitation could be addressed by integrating them into refined computational pipelines for site localization^24^ (Figure 4a). Briefly, the workflow begins with PSMs provided by a search engine (Methods). We systematically generate all potential phosphorylation-site isomers for every proposed PSM in Oktoberfest and employ Prosit-22PTM for iRT and FII prediction. These can be used to generate data-driven features that now capture the phosphorylation-site-specific properties of peptides. We leverage these features to calculate localization probabilities (Methods). After selecting the highest-scoring PTM site for each MS/MS, intensity-based features are used to estimate false discovery rate (FDR) with Percolator^28^ or Mokapot^29^.

**Figure 4.**
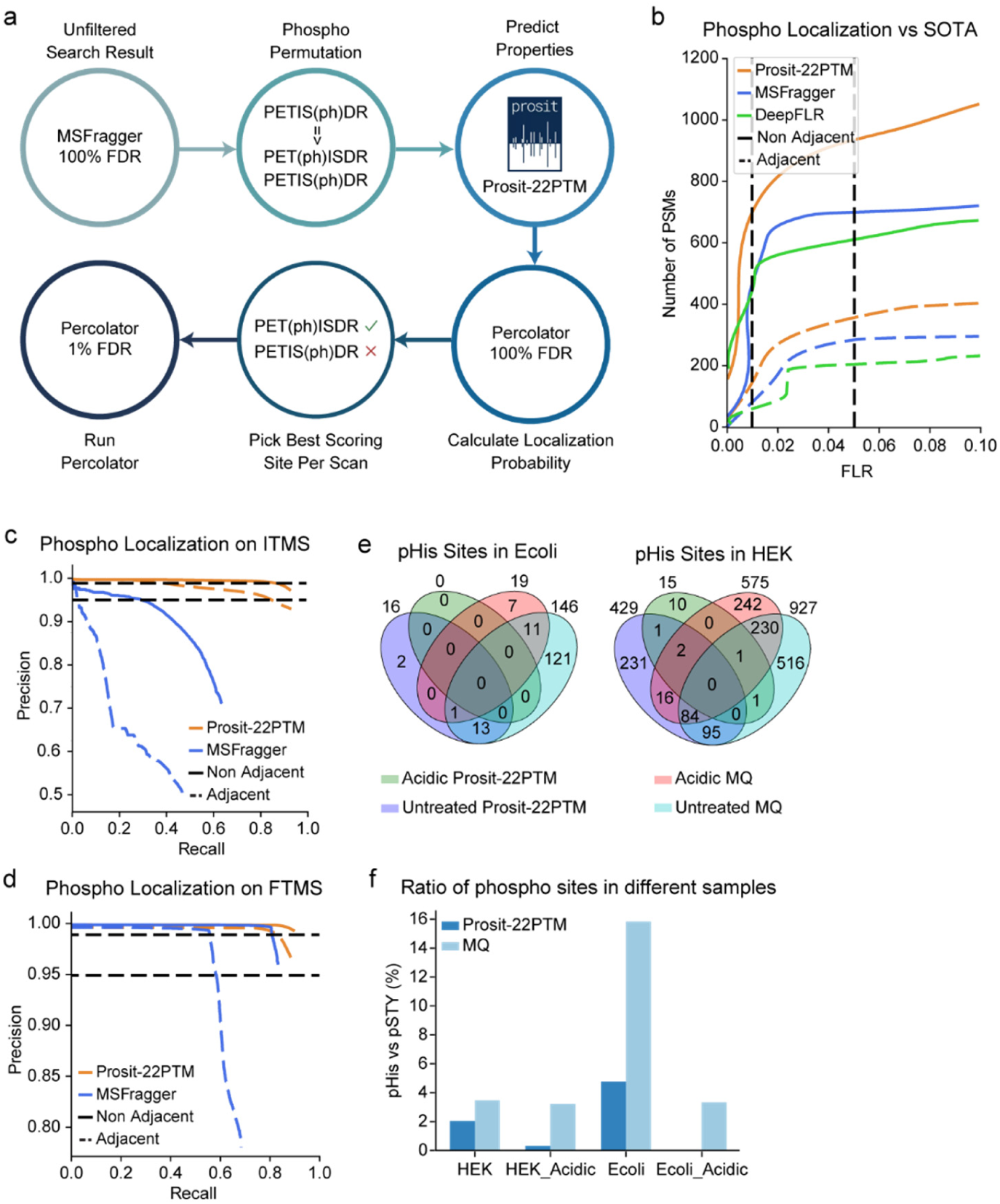
Data-Driven Phosphoproteomics Localization Pipeline for Accurate Site Assignment. **a** The localization pipeline comprises six sequential stages: initial database searching using MSFragger, in silico isomer generation through phospho permutation, scoring of all potential sites with predicted properties from Prosit-22PTM, picking the site with the highest localization score, and subsequent application of Percolator with filtering at a 1% False Discovery Rate (FDR). **b** The line plots present phosphosite localization performance. Prosit-22PTM (orange) is compared to MSFragger (blue) and DeepFLR (green) on nonadjacent (solid lines) and adjacent phospho residues (dashed lines). Vertical black dashed lines indicate 0.01 and 0.05 FLR thresholds. **c** Precision-recall curves for phosphosite localization of ITMS data. Prosit-22PTM (orange) and MSFragger (blue) performance are shown, on nonadjacent (solid lines) and adjacent phospho residues (dashed lines). Black dashed lines indicate 95% and 99% precision thresholds. **d** Precision-recall curves for phosphosite localization of FTMS data. Prosit-22PTM (orange) and MSFragger (blue) performance are shown, on nonadjacent (solid lines) and adjacent phospho residues (dashed lines). Black dashed lines indicate 95% and 99% precision thresholds. **e** Venn diagrams show the distribution and overlap of putative phosphohistidine (pH) sites in *E. coli* (left) and HEK293 cells (right) under different experimental conditions (untreated, acid-treated) and computational workflows (Prosit-22PTM in purple/green, MaxQuant in light blue/pink). **f** Bar graph shows the ratio of phosphohistidine (pH) sites to phospho-serine/threonine/tyrosine (pSTY) sites as a percentage across different biological samples and analytical conditions.

To evaluate the pipeline’s performance, we reanalyzed a previously published dataset of synthetic phosphopeptides^30^. First, we compared sorting PSMs by Percolator score versus localization probability. While both yield the same overall precision, we observe that localization probability provides higher recall (0.92 vs 0.27) at 0.99 precision (Supplementary Figure S6a). The localization pipeline also demonstrates marked improvements over existing methods, increasing the number of confidently matched PSMs by 36% at a 0.05 false localization rate (FLR) compared to MSFragger and by 52% compared to DeepFLR^31^ (Figure 4b). For sites with adjacent residues capable of carrying phosphorylation, Prosit-22PTM maintains 350 PSMs at an FLR of 0.05, which is an increase of 25% compared to MSFragger’s 280 PSMs and 75% compared to DeepFLR’s 200 PSMs.

To validate the pipeline more systematically, we evaluated it on a synthetic peptide set in which all possible amino acid substitutions were synthesized at phosphorylated positions, with a phosphorylated serine systematically replaced by threonine or tyrosine in either its phosphorylated or unmodified form. This design created a comprehensive benchmark set that extended beyond simple site permutations, enabling rigorous assessment of the pipeline’s ability to discriminate between closely related phospho-variants and their non–phosphorylated counterparts. Recall was calculated against the union of correct PSMs identified by MSFragger and Prosit-22PTM. In this benchmark, Prosit-22PTM maintained a precision above 95% up to a recall of 0.85 on spectra acquired with the Iontrap, whereas MSFragger showed a sharp decline beyond a recall of 0.4 (Figure 4c). On Orbitrap data, Prosit-22PTM showed an improvement, achieving 0.9 recall at 99% precision, compared to 0.82 recall for MSFragger. Prosit-22PTM demonstrated a significant improvement when considering only sites with adjacent residues capable of carrying phospho, with a 0.84 recall at 99% precision, compared to 0.6 with MSFragger (Figure 4d). In addition, residue-specific analyses showed that our pipeline maintained a recall of ∼0.9 at 95% precision across pS, pT, and pY, whereas MSFragger displayed a marked bias toward pY with ∼0.9 recall, but substantially lower recall of ∼0.6 for pT and pS (Supplementary Figure S6e–g).

With the ability to confidently localize phosphorylation sites, we next explored whether the model could shed light on histidine phosphorylation, given its zero-shot prediction capabilities. For this, we re-analyzed a publicly available dataset^32^, which examined the effect of acid on phosphorylated histidines (pHis). The authors argue that, as pH decreases, fewer phosphorylated histidines should be detected. Applying our pipeline to this dataset showed that *Escherichia coli* (*E. coli*) samples had 16 pH sites under untreated conditions, with no evidence for a single pHis site after acid treatment. Similarly, HEK cells had 429 sites in untreated conditions compared to 15 sites in acidic conditions, highlighting the high acid-lability of pHis in both organisms (Figure 4e). Notably, most sites originally assigned by MaxQuant (MQ) to histidine were reclassified by our pipeline as serines, threonines, or tyrosines, explaining much of the decrease in phosphohistidine site counts after rescoring. The phosphohistidine-to-canonical phosphosite (S, T, Y) ratio decreased sharply upon acidic treatment, dropping by 10-fold in HEK cells and reaching 0% in *E. Coli*, compared to ratios for untreated samples of 2.2% and 15%, respectively (Figure 4f). These results align with the hypothesis from the original study^32^, which shows a sharp reduction in phospho histidines under acidic conditions in both HEK cell lines and *E. Coli*.

By combining high-precision localization with an integrated rescoring step, our method not only enhances accuracy but also increases recall. This integrated workflow enables precise detection and annotation of challenging PTM sites across a broad range of experimental and synthetic datasets, providing a flexible framework for phosphosite analysis.

### Advancing Histone Proteomics with Machine Learning-Driven PTM Analysis

The study of multiply modified peptides remains a particular challenge due to the cumulative effects that each PTM has on FII and iRT. To evaluate Prosit-22PTMs performance on multiply modified peptides, we generated and analyzed a comprehensive panel of synthetic peptides that feature diverse combinations of 7 PTMs, with some peptides carrying at most 4 PTMs (Method) that were not used during training. First, we assessed our predictions for iRT and found that Prosit-22PTM delivers accurate predictions across all 45 tested PTM combinations, with median absolute RT errors less than 2 minutes on 60 minutes gradient (Figure 5a). Regarding FII prediction, we consistently observe median SAs exceeding 0.85 (Figure 5b). However, some combinations perform worse (e.g., peptide carrying phosphorylation, ubiquitination, and citrullination) than others which we attribute to the limited number of spectra available for these specific combinations and the complexity of the resulting charge distribution of some ions. We compared our model to state-of-the-art methods for both iRT and FII prediction and found that it consistently outperforms existing approaches across all modification levels (Supplementary Figure S7).

**Figure 5.**
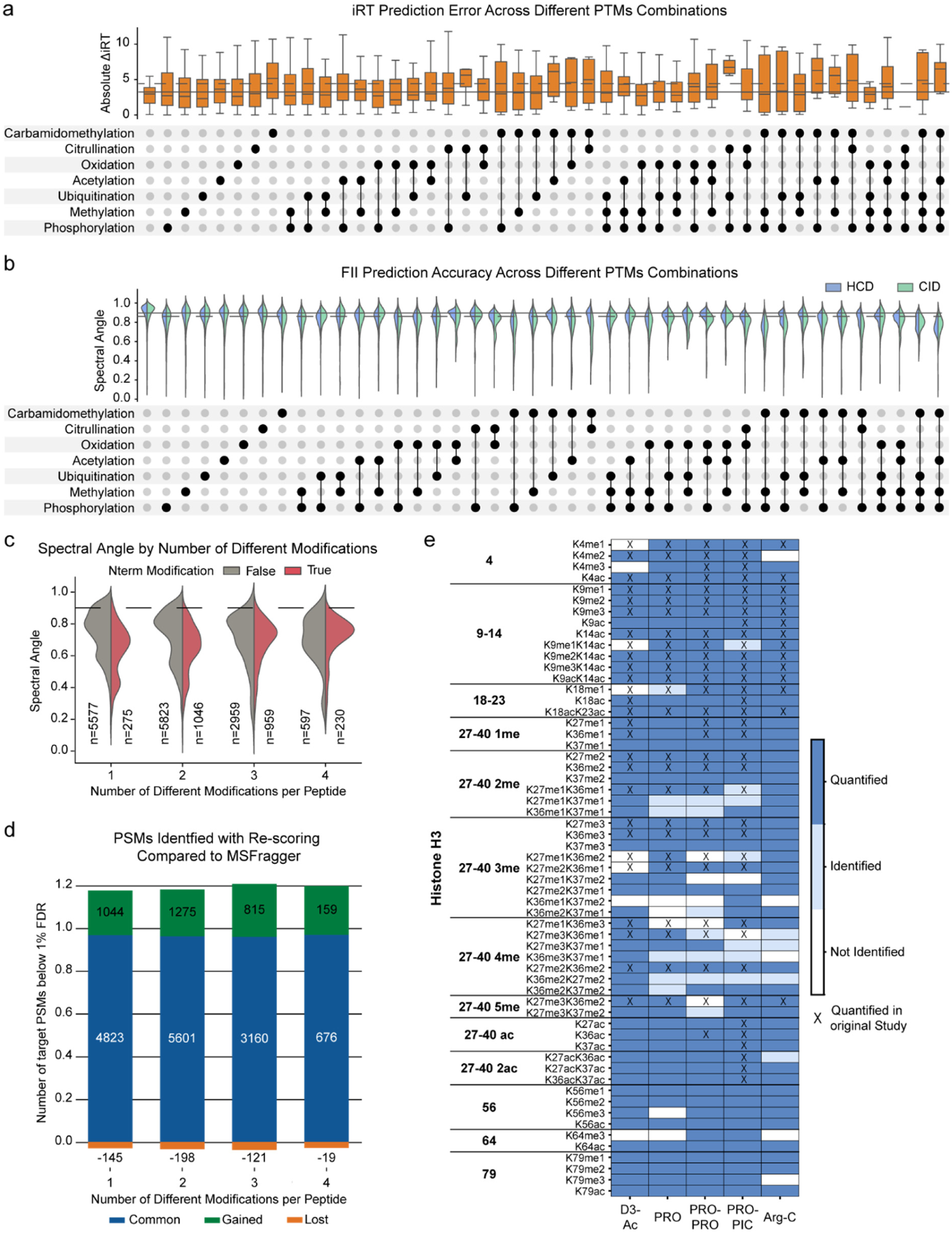
Accurate Prediction Boosts Histone Peptide Identification with Multiple Post-Translational Modifications. **a** iRT prediction accuracy with Prosit-22PTM model across various PTM combinations. The upper panel displays box plots of absolute delta iRT values. The lower UpSet plot shows the specific PTM combinations in each group, with filled black circles denoting the presence of a modification and connecting lines indicating multiply modified peptides. Solid line shows the average performance on seen PTMs, and the dashed line on unseen PTMs. **b** FII prediction accuracy with Prosit-22PTM model across various PTM combinations. The upper panel shows violin plots of spectral angle distributions. The lower UpSet plot illustrates the specific PTM combinations in each group, with filled black circles indicating the presence of a modification and connecting lines showing multiply modified peptides. The solid line represents the average performance on observed PTMs, while the dashed line indicates performance on unobserved PTMs. **c** Violin plot shows the distribution of spectral angles between predicted and experimental MS/MS spectra for peptides with 1-4 different PTMs. Peptides are grouped by the presence (red) or absence (grey) of N-terminal modifications. **d** Comparison of PSMs identified below 1% FDR before and after rescoring with Prosit-22PTM predictions. Bars show the number of target PSMs for peptides with 1-4 distinct modifications. Blue indicates PSMs identified by MSFragger and rescoring, green shows additional PSMs gained after rescoring, and orange represents PSMs lost following rescoring. **e** Histone PTM identification and quantification performance across different sample preparation methods after rescoring. The heatmap shows detection results for modified peptides organized by amino acid position and modification type. Columns represent various sample preparation protocols. Dark blue indicates peptides that were both identified and quantified, medium blue shows peptides that were only identified, and white signifies peptides that were not identified. Boxes marked with X denote peptides quantified in the original study.

Building on this foundation, we applied rescoring using Prosit-22PTM to peptides from histones with heavily modified tails from a previously published dataset^33^. Prosit-22PTM demonstrated strong FII prediction accuracy for all peptides, consistently maintaining spectral angles above 0.8 (Figure 5c), regardless of the number of modifications. However, we noted a decline in performance for peptides with N-terminal PTMs, which can be partly attributed to the limited diversity of these modifications in the training dataset. Still, applying rescoring with Oktoberfest^19^ led to a ∼20% increase in confidently identified PSMs across different modification counts compared to MSFragger (Figure 5d).

Finally, we assessed the impact of various sample preparation strategies on the identification and quantification of PTMs after rescoring. In the original study^33^, the authors evaluated various in-gel digestion and labeling strategies and found that the most effective protocol, involving Lysine and N–terminus labeling, yielded the highest number of histone-modified peptide identifications. Using our rescoring pipeline (Methods), we found that both the simpler in-solution Arg-C digest and the most complex workflow (PRO-PIC) yielded 54 quantified peptides (Figure 5e). Furthermore, we identified and quantified additional isobaric Histone H3 peptides and known sites on Lysine 56^34–36^ and 64^37,38^ that were not identified in the original study.

This analysis shows that Prosit-22PTM is a valuable tool for analyzing multiply modified peptides. Our prediction pipeline can replace complex lab workflows, simplifying and unifying processes and achieving consistent results.

### Prosit-22PTM Embeddings Reveal Physicochemical Parallels Between Modified and Unmodified Residues in HLA Peptides

To further examine the zero-shot capabilities of Prosit-22PTM, we expand our analysis from known PTM-residue combinations and multiple modifications to unseen PTMs on experimental data. This aims to assess the model’s ability to generalize beyond what was seen in the training data. For this purpose, we reanalyzed various HLA datasets^39^ searching for 34 different residue-PTM combinations which allows to evaluate three types of PTM: first, 11 AA-PTM combinations that were included in the training set that we will refer to as seen AA seen PTM combinations, second, 15 combinations where a PTM was included in the training set but on different residues (e.g. methylation on D) that we will refer to as unseen AA–seen PTM combinations, and lastly 8 combinations that were not included in the training set at all (e.g. sumoylation) that we will refer to as unseen AA–unseen PTM combinations (Methods). After rescoring using the Prosit-22PTM model, we observed a 70% increase in confidently identified PSMs. The additional identifications spread across various PTMs, regardless of whether the AA-PTM combination was observed during training, with no evident bias towards PTMs previously not seen (Figure 6a). The exception was FAT10, which exhibited a 45% reduction in PSMs.

**Figure 6.**
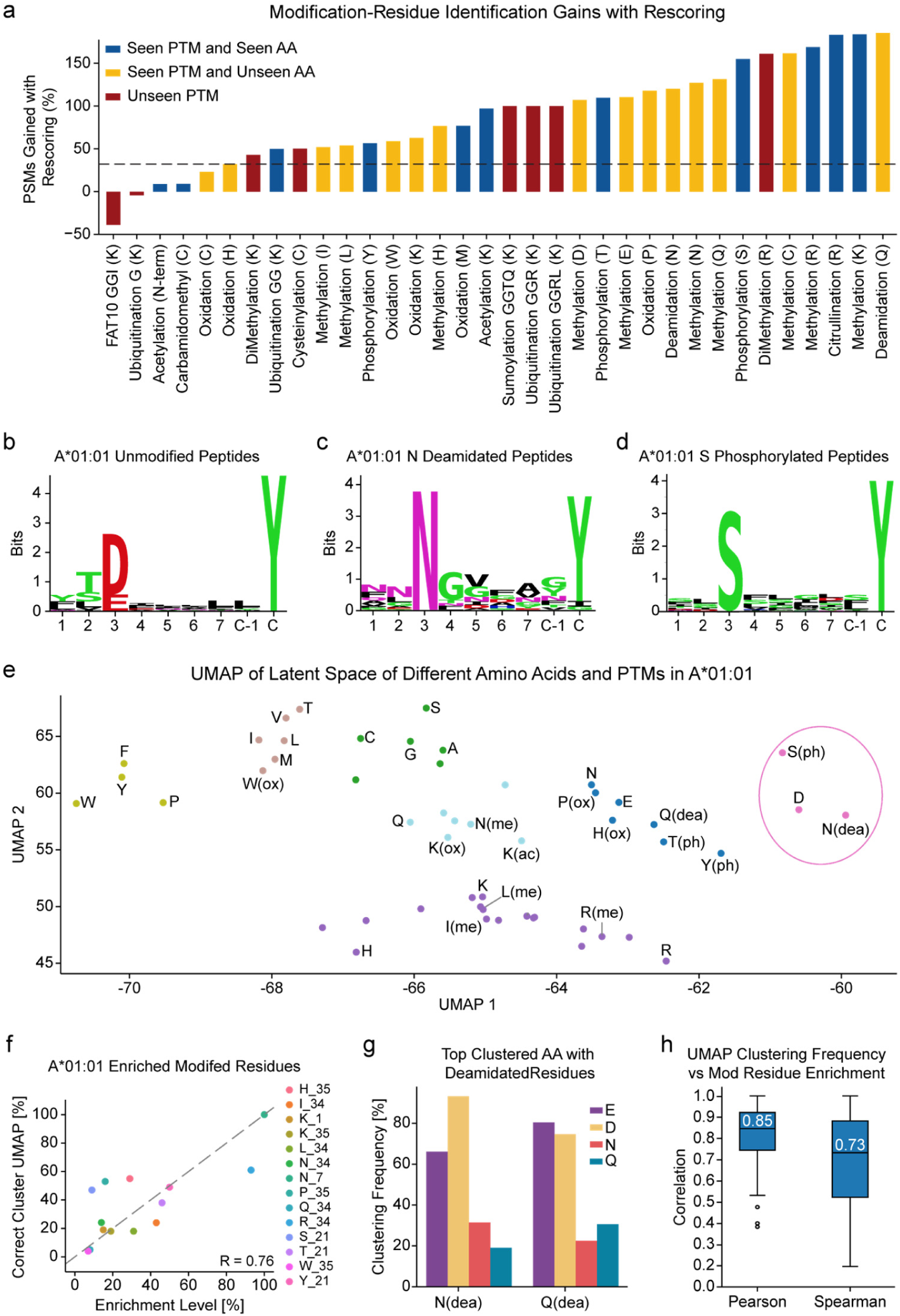
Enhanced Detection and Motif Extraction for Modified HLA Peptides. **a** Bar plot shows the increase in PSMs after rescoring HLA datasets with Prosit-22PTM for different modification-residue combinations. Bars are colored by category: blue for seen PTMs, yellow for unseen modification-residue combinations, and red for unseen PTMs. **b** Peptide motif plot for the unmodified peptides in HLA-A*01:01.The color coding follows standard AA physio-chemical properties (black hydrophobic, red acidic, blue basic, purple neutral, and green polar). **c** Peptide motif plot for the peptides with deamidated Asparagine in A*01:01. **d** Peptide motif plot for the peptides with phosphorylated Serine in A*01:01. **e** UMAP projection of latent space embeddings from Prosit-22PTM, highlighting relationships between unmodified and modified AAs. The UMAP was trained on the embeddings of unmodified AAs (black labels). **f** The scatter plot shows that Modification enrichment in A*01:01-presented peptides correlates with clustering in UMAP embeddings across 100 runs with varied parameters. Enrichment levels are normalized (x-axis, with the highest set to 100%) against the percentage of UMAP runs where modified residues clustered with unmodified AAs in the same binding position (y-axis). Each point represents an AA modification (legend: single letter code + unimodal id). **g** The grouped bar chart displays the clustering frequency of deamidated residues with unmodified AAs across UMAP projections for all 92 HLA alleles analyzed. Y-axis shows the percentage of UMAP runs where deamidated asparagine [N(dea)] and deamidated glutamine [Q(dea)] clustered with each of the four AAs: (E, purple), (D, orange), (N, red), and (Q, teal). **h** Box plots comparing Pearson and Spearman correlation coefficients between UMAP clustering frequency and modified residue enrichment levels across all 92 HLA alleles. Each data point within the distribution represents a single HLA allele, quantifying the extent to which the relationship between modification enrichment and correct clustering holds for that specific allele.

Next, we wanted to assess if the additional peptides identified display the anticipated binding motifs. To evaluate whether the detected peptide-binding motifs, both canonical and modified, align with established literature, we focused on analyzing HLA class I peptides derived from 95 HLA monoallelic cell lines^40^ where the expected motifs are well described. As an example, we extracted unmodified peptide motifs associated with allele A*01:01 from the rescored results (Figure 6b), in which we observe a preference for aspartic acid (D) at position 3, consistent with motifs reported in previous immunopeptidomic studies^10,39^. Extending our analysis to modified peptides revealed patterns consistent with the physicochemical properties of amino acids. Deamidated asparagine (N[dea]) modified peptides bound to HLA-A*01:01 showed a significant enrichment of modified N at position 3 (Figure 6c), mirroring the unmodified motif, as this position now contains the deamidation product, which is chemically identical to aspartic acid residue. Similarly, phosphoserine (S[ph]) modified peptides exhibited enrichment patterns in which S appeared to replace aspartic acid at position 3 (Figure 6d), consistent with previous studies^41,42^.

Since the substitution patterns correlated with the physicochemical properties of the modified residues, we hypothesized that the representations learned by Prosit-22PTM could encapsulate the HLA binding preferences for modified and unmodified residues within the embeddings. To explore this hypothesis, we extracted embeddings from the model’s internal representations and utilized semi-supervised UMAP dimensionality reduction to visualize the organization of unmodified and modified AAs for each allele separately (Methods). The resulting embedding space for A*01:01 displayed clear clustering; for instance, phosphoserine and deamidated arginine formed a cluster alongside aspartic acid (Figure 6e), reflecting the similarities noted in the binding motifs without access to the list of identified peptides. Additionally, we observed other enriched modifications associated with A*01:01 that grouped closely with glutamic acid (E), which is also prevalent at position 3. Notably, deamidated glutamine (Q[dea]), another enriched PTM in position 3, is clustered together with E. Phosphorylated threonine (T[ph]) is also clustered with E, consistent with the established use of E as a proxy for T[ph] in HLA motif predictions^41,42^.

To assess whether this physicochemical organization was consistent beyond isolated cases, we systematically evaluated clustering stability across 100 independent UMAP runs (Methods). In the A*01:01 allele, higher enrichment levels of modified residues were strongly associated with the proportion of correctly clustered modified amino acids at anchor positions (R=0.76) (Figure 6f). This indicates that biologically relevant modifications—those most common in HLA-presented peptides—were clustered more often with their physicochemical counterparts. This pattern suggests it is applicable across the broader HLA landscape. Examination of deamidation clustering among all 95 HLA alleles reveals a consistent physicochemical logic: deamidated asparagine associates with aspartic acid in 90% of the cases, and deamidated glutamine aligns with glutamic acid in 80% of runs (Figure 6g). The robustness of the relationship between enrichment and clustering across various HLA alleles is notable, with median Pearson correlations of 0.85 and Spearman correlations of 0.73 between modification enrichment and correct clustering frequency (Figure 6h).

The systematic relationship observed here indicates that the latent representations in Prosit-22PTM may also effectively encode core physicochemical principles beyond peptide fragmentation. The organized structure of PTMs within the model’s latent space opens promising avenues for computational research in modified peptide analysis, exemplified by motif prediction tailored to specific HLA alleles.

### Achieving Robust PTM Prediction Solely Through Amino Acid Substitution Augmentation

With the success of data augmentation, we hypothesized that this strategy can be applied to enhance any training dataset, particularly those that lack substantial PTM examples, enabling the learning and prediction of peptide properties for modified peptides. To test this, we trained models with augmentation (WA; implied when omitted) and without augmentation (WOA; explicitly stated if trained without augmentation) using the previously published ProteomeTools data exclusively (Prosit-2PTM), meaning, absent the PTM extension introduced here, and using all now available ProteomeTools data (Prosit-22PTM). Prosit-2PTM-WA still demonstrated strong signs of accurate zero-shot predictions, despite lacking any relevant PTM data for training, with an average SA of 0.84 across all modified peptides compared to 0.71 and 0.87 for Prosit-2PTM-WA and Prosit-22PTM-WOA, respectively (Supplementary Figure S8a). This is particularly evident on biotinylation where Prosit-2PTM-WA reaches a prediction performance of 0.81 SA compared to 0.56 for Prosit-22PTM-WOA. However, Prosit-2PTM-WA performed poorly on phosphorylated peptides only reaching an SA of 0.75 compared to 0.92 for Prosit-22PTM-WOA, due to the lack of phosphorus atoms in the augmentations. Despite this, AAS-based augmentation is a highly valuable strategy for substantially improving model performance beyond what is available for training and can lead to good generalization. This is particularly relevant when training data is sparse, i.e., for newly released mass spectrometers and understudied fragmentation methods, and likely applies to most peptide properties, where (extensive) PTM training may not yet be available.

## Discussion

This study introduces ProteomeTools-PTMs, a comprehensive and openly available resource that accelerates the study of post-translational modifications (PTMs) in proteomics. Importantly, the dataset’s scale and diversity are expanded through a data augmentation strategy that enables robust model generalization, especially for unseen or rare PTMs. This advancement is not feasible with chemical synthesis alone due to the 10-fold increase in chemical space achieved through augmentation. Using this augmented PTM data, we developed Prosit-PTM, a deep learning model for predicting indexed retention times and fragment ion intensities for modified peptides. Prosit-PTM delivers significant improvements in accuracy and generalization, especially for modifications not seen during training in a zero-shot manner, surpassing current state-of-the-art methods^17,18^. This model, using large-scale augmentation, sets a new standard for PTM-aware predictions.

Our results demonstrate performance improvements in phospho localization (recall reaching up to 90% at less than 1% false localization rate) and a consistent 20% increase in the identification of multiply modified peptides, highlighting immediate benefits for high-confidence PTM assignments. Furthermore, the model’s ability to encode detailed representations of modifications and amino acids similar to foundational models opens exciting possibilities for other amino acid property prediction models, showcased on PTM-aware HLA-binding prediction models, potentially advancing precision immunopeptidomics.

All the data, models, and pipelines used in this publication including the physical synthetic peptides are available to the community free-of-charge, supporting various user groups. First, clones of the synthetic peptide libraries are available upon request, enabling the characterization of newly developed mass spectrometers and algorithms^43^. Second, all raw data acquired in the ProteomeTools project are available on PRIDE and are used frequently for model development and algorithm validation^44^. Third, the source code for model training (https://github.com/wilhelm-lab/dlomix), online model inference through Koina^45^, and rescoring, including site localization and spectral library generation, through Oktoberfest^19^ will assist model developers, software developers, and wet-lab scientists.

Although not demonstrated in this work, Prosit-PTM can be used to validate open PTM search results by predicting MS2 spectra and retention times for a wide range of modified peptides, including those with unknown or novel PTMs. Also, the augmentation strategy developed here is likely applicable to other fields predicting properties of molecules beyond peptides as the augmentation can be expanded to capture chemical reactions, rather than amino acid substitutions. In summary, this work marks a crucial step toward improving the rigor of PTM validation, leading to deeper insights into PTM dynamics, crosstalk, and regulation and we further anticipate that Prosit-PTM will be of benefit for the scientific community with impact beyond proteomics research.

## Methods

### Proteometools

#### Peptide Sets

Peptides selected for synthesis can be organized into 33 distinct sets based on the PTM included in the set. Initially, we developed sets that each focus on a single unique post-translational modification (PTM), including Acetylation, Citrullination, Monomethylation (on arginine and lysine), O-GalNAc (on serine and threonine), O-GlcNAc (on serine and threonine), and Phosphorylation (which includes three sets, one for each of serine, threonine, and tyrosine). We also labeled some of this with TMT-6plex, specifically: Acetylation, Monomethylation, Phosphorylation, and Ubiquitination. Furthermore we labeled 4 different tryptic peptides^9,22^ namely first pool, second pool, isoform and thermo_SRM with di-methylation and acquired in CID and HCD fragmentation methods. Additionally, we explored other specialized sets to validate various downstream analyses. The first specialized set consisted of previously annotated human phospho-sites from the PhosphoSitePlus database to validate various phospho-localization pipelines. We established additional sets that included permuted phosphorylation across various residues eligible for modification. The permutation involved relocating the phosphorylation to different positions within the peptide while remaining on the same residue, resulting in three distinct sets—one for each of the STY amino acids. These sets can also be used to validate diverse phospho-localization pipelines. Lastly, our specialized set, combPTM, included multiply modified peptides containing 7 unique post-translational modifications (PTMs), with each peptide allowing up to 4 modifications. In total, we generated 42 unique combinations of PTMs; this set aimed to replicate the complex effects of multiply modified peptides typically found in histones. The methodologies for designing peptide pools, synthesizing peptides, preparing samples, and analyzing synthetic peptides using liquid chromatography-tandem mass spectrometry (LC-MS) have been comprehensively described in the first Proteometools publication^22^.

#### Quality Control Peptides

Each sample was spiked with two retention time standards (Pierce Retention Time Standard and PROCAL)^46^. These peptides were used to calculate iRT, which reduces retention-time variability across different acquisitions^22^. Peptide sequence TFAHTESHISK was picked as the left anchor with value 0, and SILDYVSLVEK was picked as the right anchor with value 100. These two peptides were selected based on their low variance and their identification in the majority of experiments. When one of these peptides was not found, the peptide with the closest retention time was selected, along with its respective iRT. For aligning MS2 spectra across different measurements, the same set of QC peptides was used to calibrate the collision energy. The alignment was performed as described in the PROCAL publication^46^.

#### Data Acquisition

Data acquisition was performed as previously described^10^. Briefly, we utilized five different fragmentation methods, including the most commonly used ones like collision-induced dissociation (CID; also known as resonance-type CID) and higher-energy CID (HCD; also referred to as beam-type collision-induced dissociation), with six collision energies (23, 25, 28, 30, 35) for HCD fragmentation. CID was exclusively acquired on an Ion Trap Mass Analyzer, while most of the HCD data was collected on an Orbitrap; only one collision energy (35) was obtained with the Ion Trap. Electron-transfer dissociation (ETD) was also used, involving an ETD and FTMS scan (charge-dependent reaction time), electron-transfer/collision-induced dissociation (EtciD) with NCE 35 FTMS, and electron-transfer/HCD (EthcD) with NCE 28 FTMS.

#### Database searching

Raw data acquired were analyzed using MaxQuant v.1.5.3.30, with individual LC–MS runs searched against pool-specific databases. The first search tolerance was set to 20 ppm, and the main search tolerance was at 4.5 ppm, with filtering for PSM and protein FDR at 1%. Methionine oxidation was included as a variable modification in all searches; other fixed and variable modifications depended on the PTMs present in each set.

### Model Data Preparation and Training

#### Data preparation

Results from MaxQuant were annotated according to the assigned peptides. For all PTMs, if MaxQuant misassigned the site, this was corrected and transferred to the site based on the expected synthesized sequence. All resulting PSMs were filtered with an Andromeda score greater than 70; any PSMs scoring below 70 were discarded. For FII data, canonical y- and b-ions were annotated with charges from 1 to 3 and ions from 1 to 29. Precursor ions were annotated, and peaks were removed to ensure they were not assigned to any other ion. Intensities were base peak normalized so that the highest annotated ion is assigned a value of 1, and all other ions are scaled between 0 and 1 depending on their ratio to the base peak. PSMs were grouped by modified peptide sequence, precursor charge, and calibrated collision energy, with a maximum of 3 PSMs selected per group. After these steps, 30.4 million PSMs remained and were used for model development. For IRT data, PSMs were grouped by modified peptide sequence, and peptides with variance less than 2 iRT units were selected. For the peptides chosen, the mean iRT value was used as the prediction label. After these steps, 1.5 million peptides remained and were used for model development.

#### Data Augmentation

We incorporated all amino acid (AA) substitutions (AAS) listed in the Unimod database (https://unimod.org/) to thoroughly enrich our dataset. Each substitution was categorized into six distinct classes based on its effects on mass-to-charge ratios (MZ). This classification was consistently applied, taking into account the influence of these substitutions on MZ. Additionally, we calculated the delta iRT 95 and spectral angle based on the differences between the original AA and its substituted form, subsequently classifying these into six categories to reflect their impacts. To ensure our datasets comprising training, validation, and test sets contained a comprehensive array of potential combinations, we used the IterativeStratification function from the scikit-learn library. This method allowed us to achieve balanced representation across the various class combinations within each subset. We also ensured that each substitution and its reverse were included in the same set. By maintaining this balance, we aimed to minimize bias in our evaluation processes while ensuring that all classifications were adequately represented in the analyses. After splitting, we ended up with 258 AAS in the train set, 42 in the val set, and 42 in the test set. Because of combinatorics and the wide variety of available AAS, we can scale any training data to nearly unlimited size. For practical reasons, we limited augmentation to a maximum of one additional modified sequence per spectrum. To ensure adequate coverage of all available AAS and the chemical space, we sampled a single AAS per original training sample uniformly (Supplementary Figure S5c). To expand our coverage of chemical space and enable the generalization of more complex PTMs, we also applied AAS to residues that were already modified (e.g., S[ph] -> A[A→S][ph]).

#### Data Splitting

We split the data into training, validation, and test sets, ensuring that the splits were based on base peptide sequences^47^. This approach allows for various modified versions of the same peptide sequence to remain within the same set. To prevent data leakage with AAs, substitutions (e.g., A[Ala→Met]) and their reverse (M[Met→Ala]) for one peptide are always in the same set—training, validation, or holdout—or are not generated at all when the respective base sequences are assigned to different sets (e.g., if PEPTIDEK is in training and PTPTIDEK is in validation, then PT[T->E]PTIDEK would not be generated). To effectively evaluate our models against previously unseen post-translational modifications (PTMs), we intentionally created multiple holdout sets, as recommended by ProspectPTMs^48^. Additionally, we excluded all sequences from our training and validation sets that correspond to peptide sequences previously published in ProteomeTools 21PTMs^23^. We established two distinct testing sets: one comprising unseen peptide sequences paired with seen residue-PTM combinations, and another consisting of both unseen peptide sequences and unseen residue-PTM combinations. This approach facilitates the assessment of the model’s ability to generalize to previously unencountered peptides and to new residue-PTM combinations. Following data partitioning for the development of the FII model, we compiled a training dataset comprising 55 million PSMs, along with 14.4 million PSMs for validation and 7.3 million PSMs for testing. For the development of the iRT model, we organized our datasets similarly, resulting in 2.7 million peptides for training, 684,000 peptides for validation, and 346,000 peptides for testing.

#### PTM Encoding

We utilized Unimod (https://unimod.org/) to extract delta mass and atom count for PTM encodings. For gain and loss encodings, we calculated this based on the closest carbon atom to where the chemical structure is modified. The delta of gain minus loss is equal to the delta atom count introduced by the PTM. In certain cases, we account for carbon atoms in the loss, even when there is no actual addition or removal of carbon atoms from the molecule; this occurs when a modification involves a rearrangement of existing functional groups, such as the conversion of an amine to a carbonyl group or the formation of a cyclic structure. One of the modifications that leads to this is the change of glutamic acid to pyroglutamic acid, which results in a cyclic carbon structure that does not exist in unmodified glutamic acid. To accurately encode these changes, we represent the transformation as a loss of carbon from the original functional group and a gain of carbon in the new functional group, even though the total number of carbon atoms remains unchanged. This approach enables us to simulate the chemical changes while ensuring that the net delta (gain minus loss) accurately reflects the change in atom count introduced by the PTM. This method maintains consistency with Unimod’s mass and atom accounting while capturing the nuanced structural rearrangements that occur.

#### Model Architecture

Both iRT and FII models were implemented using Keras with TensorFlow 2.x as the backend and share a common sequence encoder architecture designed to process peptide sequences alongside their position-specific post-translational modification (PTM) features. Each peptide, padded to a fixed length of 32 to include N- and C-termini, is integer-encoded and passed through an embedding layer that produces a (32, 16) representation. PTM features corresponding to each residue are processed through a dedicated fully connected network consisting of three dense layers with 1,024, 64, and 16 units, each followed by a dropout rate of 0.3. The encoded PTM vector is concatenated with the sequence embedding, resulting in a fused peptide representation of shape (32, 32). This fused input is passed through a bidirectional GRU layer with 256 units and a subsequent GRU layer with 512 units, both followed by dropout (rate 0.3). The encoded sequence is then transformed via a global attention mechanism that yields a contextual vector of size 1,024, which serves as the encoded representation shared between the two Prosit models.

For the iRT model, the shared encoder output is processed by a dense regression head of 512 units with ReLU activation and a dropout rate of 0.1, followed by a final dense layer with a single unit that predicts the indexed retention time value. This architecture allows both models to leverage identical sequence and PTM encoders while diverging only at their respective decoding stages to suit the distinct regression tasks of retention time and spectral intensity prediction.

For the FII model, an additional metadata encoder is incorporated to capture experimental parameters such as precursor charge, fragmentation type, and collision energy. These variables are processed through a dense layer of 512 units followed by dropout (rate 0.3). The resulting metadata embedding is broadcast to match the sequence context and fused with the sequence-PTM representation using element-wise multiplication. The combined representation is then passed through an attention layer that integrates information across sequence positions and PTM sites. Decoding is performed using a GRU layer with 512 units followed by dropout (rate 0.5) and a decoder-specific attention mechanism, yielding position-specific fragment context vectors. The output passes through a time-distributed dense regressor with six units and leaky ReLU activation, which is flattened into a 174-dimensional vector corresponding to predicted fragment ion intensities.

#### Model Training

For the iRT model, mean absolute error was used as the loss function. The initial learning rate was 0.001, with a reduction factor of 0.8 when the validation loss did not improve over 5 epochs, and 0.00001 was set as the minimum learning rate. The model trained for up to 160 epochs with early stopping if the validation loss did not improve for 10 epochs. For the FII model, normalized spectral contrast loss^9^ was used as the loss function. We employed the Adam optimizer with a cyclic learning rate (CLR) algorithm^49^. During training, the learning rate cycled between a fixed lower limit (0.00001) and an upper limit (0.0002), scaled by a factor of 0.95 every 8 epochs. The model trained for a maximum of 120 epochs with early stopping if the validation loss did not improve over 16 epochs.

### Application and Calibration of State-of-the-Art Models

#### Koina

Predictions were generated using Koina^45^, via Python-based Koinapy package. This enabled streamlined, programmatic access to get predictions from DeepLC and AlphaPeptDeep models. For each raw file, relevant peptide sequences and associated metadata (such as precursor charge and collision energy) were submitted to Koina, and the resulting predictions were retrieved for downstream processing.

#### DeepLC

Retention time (RT) predictions were generated using DeepLC^17^. To ensure these predictions accurately reflected the experimental setup, an iRT (indexed Retention Time) calibration was performed individually for each raw file. This involved using a set of unmodified peptides with known experimental retention times, which were present in each file. Predicted RTs were regressed against observed iRT values, and the resulting calibration was applied to all DeepLC predictions within that raw file, correcting for systematic deviations and aligning predictions to the experimental RT scale.

#### AlphaPeptDeep

For intensity prediction, AlphaPeptDeep^18^ was used. To account for instrument-specific fragmentation, collision energy (CE) calibration was performed per raw file. This process involved adjusting the CE parameter based on the set of unmodified peptides in each raw file to obtain the best-fitting calibrated CE that leads to the highest spectral angle. The calibrated CE value was then applied to all AlphaPeptDeep predictions for that file, ensuring that intensity predictions were tailored to the specific experimental conditions of each run.

### General Rescoring and Localization Pipeline

#### Rescoring Pipeline

To ensure accurate false discovery rate (FDR) estimation and filtering for both modified and unmodified PSMs, we implemented a tailored rescoring pipeline using Percolator. For unmodified PSMs, Percolator was run once to rescore and filter identifications using standard FDR estimation. For modified PSMs, a two-stage Percolator workflow was employed^24^. In the first stage, Percolator was run on a combined set of PSMs, where modified PSMs comprised the relevant set and their complementary PSMs. The neighbor set, formed by permuted PTMs at all possible sites, following the subset-neighbor search (SNS) approach^50^. After this run, only the highest-scoring PSM for each scan was retained. In the second stage, Percolator was run exclusively on the selected, modified PSMs per scan from the first run, using a procedure that followed the group FDR approach^51^, with a 1% FDR threshold applied at the PSM level. This two-step rescoring and filtering process ensures robust and accurate FDR control for both unmodified and modified PSMs, thereby enhancing the reliability of downstream analyses.

#### Site Localization Probability Calculation

To assess the confidence of phosphorylation site localization within peptides, we adopted a probability-based approach that leverages the posterior error probability (PEP) for each candidate site. The PEP represents the probability that a specific site assignment is incorrect, given the observed data and the scoring model. As recommended in recent literature, PEP can be used to estimate local False Localization Rate (FLR), assuming the PSM itself is correct^52^, which we ensure by applying PEP to PSMs after 1% FDR filtering. For each peptide containing multiple potential phosphorylation sites, we first calculated a site score for each candidate site as the inverse of its PEP (1/𝑃𝐸𝑃). This transformation ensures that sites with lower error probabilities (i.e., higher confidence) contribute more strongly to the final localization probability. This approach is mathematically consistent with the scoring framework described in phosphoRS^53^, in which the site score is defined as the inverse of the site-specific p-value (i.e., error probability).

To obtain a normalized probability for each site, we summed the scores of all possible phosphosite localizations within a given peptide. Then we divided the score for each site by this sum. Formally, for a peptide with 𝑛 possible phosphosites and corresponding PEP values 𝑃𝐸𝑃_1_, 𝑃𝐸𝑃_2_,. .. , 𝑃𝐸𝑃_n_, the localization probability for a site 𝑘 is calculated as:

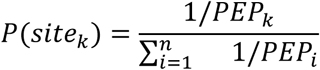

This normalization ensures that the sum of localization probabilities across all candidate sites for a peptide equals one, providing a direct and interpretable measure of confidence for each site assignment.

### Data processing

#### Synthetic Phospho Data

##### Database searching

All phospho data were analyzed using MSFragger 4.0 and FragPipe 2.12, with carbamidomethylation set as a fixed modification, and methionine oxidation, as well as phosphorylation on serine, threonine, and tyrosine, set as variable modifications, and Trypsin was used as the enzyme for digestion. Parameter calibration was enabled to calibrate raw file mz values, and the optimal MS1 and fragment ion tolerances were determined dynamically. Raw files were searched with a 1% FDR at both PSM and peptide levels with PTMProphet^54^ enabled to extract localization probabilities. For the external phospho peptides PXD007058, we used the fasta files uploaded with the original publication, and all raw files were searched together. For ProteomeTools phospho permutation data, fasta files based on peptides synthesized in all pools were used, and all raw files were searched together. For ProteomeTools phospho important data, the fasta file used was the human fasta downloaded from UniProt on 11/17/2023, containing 20,596 proteins, and all raw files were searched collectively.

##### Rescoring

The rescoring and localization pipeline outlined in the previous section is systematically applied across all datasets. PepXML files generated by FragPipe are used to extract comprehensive identifications, as they are unfiltered with a 100% FDR. All sites identified by MSFragger are subsequently re-assigned based on the localization probabilities discussed earlier. Following this, Percolator is employed exclusively on the highest-scoring site for each PSM, and the results are further refined to achieve a 1% FDR.

##### Analysis

The data were sorted by the probability scores assigned by FragPipe, DeepFLR, and the Prosit-22PTM localization pipeline. All Peptide Spectrum Matches (PSMs) that included phosphorylation were categorized into two groups: 1) Non-Adjacent: where the phosphorylation on the synthetic peptide was associated with an amino acid (AA) that does not have either S/T/Y directly adjacent to it, and 2) Adjacent: where the phosphorylation was linked to an AA that has either S/T/Y directly next to it. The False Localization Rates (FLR) were then calculated based on the synthetic peptides and the accuracy of site assignments by the various pipelines. Additionally, the same analysis was conducted for precision-recall curves.

#### Phospho Histidine

##### Database searching

All phospho data were analyzed using MSFragger 4.0 and FragPipe 2.12, with carbamidomethylation set as a fixed modification, and methionine oxidation, as well as phosphorylation on histidine, serine, threonine, and tyrosine, set as variable modifications, and Trypsin was used as the enzyme for digestion. Parameter calibration was enabled to adjust raw-file mz values, and the optimal MS1 and fragment-ion tolerances were determined dynamically. Raw files were searched with a 1% FDR at both PSM and peptide levels with PTMProphet^54^ enabled to extract localization probabilities. Either the human fasta was used or *E.coli,* depending on the raw file.

##### Rescoring

Same as previously described in Synthetic Phospho Data section.

##### FLR

Since this dataset is not synthetic, an alternative amino acid had to be selected as a decoy site. Proline was chosen as the decoy amino acid because of its high prevalence adjacent to phosphorylation sites in human cells; previous analyses indicate that approximately 40–50% of pS sites are proximal to a proline residue, furthermore, similar scoring and proximity to pS patterns have been observed for pH sites as well^32^. Additionally, permuting phosphorylation of proline residues results in approximately a twofold increase in the number of detectable sites, ensuring that both target and decoy sites have comparable numbers. The data reported in the manuscript are based on a filtering process that achieves a false localization rate (FLR) of 1% by using the number of proline sites as decoys.

#### Histones

##### Database searching

All Histone data were analyzed using MSFragger 4.0 and FragPipe 2.12, with carbamidomethylation set as a fixed modification and methionine oxidation as a variable modification, and Arg-C was used as the enzyme for digestion. In in-solution Arg-C digestions, acetylation, methylation, dimethylation, and trimethylation of Lysine were added as variable modifications. The D6-Ac and PRO protocols also incorporate D3-acetylation/propionylation (+45.0294 /+56.0262 Da), lysine monomethylation with D3-acetylation/propionylation (+59.0454/+70.0422), as well as dimethylation, trimethylation, and acetylation on Lysine as variable modifications. For the PRO-PIC and PRO-PRO protocols, propionylation, monomethylation-propionylation, dimethylation, trimethylation, and acetylation on Lysine were added as variable modifications, along with N-terminal PIC labeling (+119.0371 Da) or N-terminal propionylation (+56.0262 Da), respectively. Parameter calibration was enabled to adjust raw-file mz values, and the optimal MS1 and fragment-ion tolerances were determined dynamically. Raw files were searched with a 1% FDR at both PSM and peptide levels. Human histone fasta was used, and every experimental condition was searched separately.

##### Rescoring

Same as previously described in Synthetic Phospho Data section.

##### Analysis

To obtain quantification values after rescoring, we used MSFragger with a 100% FDR threshold. For all PSMs filtered to a 1% FDR after rescoring, we examined whether these PSMs had corresponding quantification data and mapped the quantification to each PSM. Based on the results, a PSM that appeared only in the identification results and lacked quantification was categorized as identified. If it was found in both datasets, it was labeled as quantified; otherwise, it was classified as not identified.

#### HLA

##### Database searching

All phospho data were analyzed using MSFragger 4.0 and FragPipe 2.12. The search was set to an unspecific search, with a minimum peptide length of 7 and a maximum of 14. No fixed modifications were used. Variable modifications were set to: 1-Oxidation (15.9949 da) on M, W, H, K, P, and C. 2-Acetylation (42.0106 da) on n-term and K. 3-Phosphorylation (79.96633 da) on S, T, and Y. 4-Methylation (14.01565 da) on C, H, N, Q, K, R, L, I, D, and E. 5-Di-Methylation (28.0312 da) on K and R. 6-Citrullination (0.984016 da) on R. 7-Deamidation (0.984016 da) on N and Q. 8 - Ubiqitination G (57.0215 da) on K. 9-Ubiqitination GG (114.0429 da) on K. 10-Carbamidomethylation (57.0215 da) on C. 11-Cysteinylation (119.0041 da) on C. 12-Ubiqitination GGR (270.1440 da) on K. 13-Ubiqitination GGRL (383.2281 da) on K. 14-Sumoylation GGT (215.0906 da) on K. 15-Sumoylation GGTQ (343.14917 da) on K. 16-FAT10 GGI (227.127 da) on K. 17-FAT10 GGIC (330.13617 da) on K.

Set the max variable mods per peptide to 5, with a max of 60 variable modifications to manage database size. Set the slice-db parameter to 20 to run the search without exceeding RAM usage. Raw files were searched with a 1% FDR at both PSM and peptide levels. The fasta file used was the human fasta downloaded from UniProt on 11/17/2023, containing 20,596 proteins.

##### Rescoring

Same as previously described in Synthetic Phospho Data section.

##### Motif Generation

Different HLA Alleles were searched for peptides with lengths of 8 to 11. To address length discrepancies, motifs were constructed using positions 1–7, starting from the N terminus, followed by the C terminus and its preceding position. For 9-mer epitopes, the motif is derived from all nine positions. For 8-mer peptides, the seventh position is duplicated, appearing as both positions 7 and 8/C-1. For peptides with more than 9 residues, the motif omits positions 8 through C-1. For unmodified peptides, the AAs in each position were compared with their corresponding background frequencies in humans. For modified peptides, the frequencies of the AAs from the unmodified peptides served as the background. Information content was calculated in bits and plotted using the Logomaker Python package^55^.

##### UMAP Training

In this study, we employed ParametricUMAP^56^ to cluster various residue-PTM combination embeddings alongside unmodified residues effectively. The UMAP model was trained using a comprehensive set of 100 distinct parameter combinations, with the distance metric varying from 1 to 4 in increments of 0.15 and the spread parameter ranging from 1 to 3 in increments of 0.5. Each UMAP was allele dependent where it was trained on the unmodified amino acids as anchors (e.g. in A*01:01 UMAP was given as an input that D and E in cluster 1, T and S in cluster 2 and Y alone in cluster 3). To cluster AAs and modified AAs together, HDBSCAN^57^ was used. HDBSCAN was chosen because it doesn’t require specifying the number of clusters beforehand and can identify noise as well. The only parameter provided was a minimum cluster size of 2, and each cluster was plotted in a different color based on the clustering results.

## Data availability

Annotated training data is made available on HuggingFace for indexed retention time (https://huggingface.co/datasets/Wilhelmlab/prospect-ptms-irt) and fragment ion intensities (https://huggingface.co/datasets/Wilhelmlab/prospect-ptms-ms2).

## Code availability

Source code and scripts for data preprocessing are available on GitHub via https://github.com/wilhelm-lab/PROSPECT. Source code and scripts for model training are available on GitHub via https://github.com/wilhelm-lab/dlomix. Source code and scripts for rescoring are available on GitHub via https://github.com/wilhelm-lab/oktoberfest/tree/features/ptm_pipeline. Source code and scripts for model inference are available on GitHub via https://github.com/wilhelm-lab/koina.

## Acknowledgements

Funded in part by the European Union. Views and opinions expressed are however those of the author(s) only and do not necessarily reflect those of the European Union or the European Research Council Executive Agency. Neither the European Union nor the granting authority can be held responsible for them.

This work is partly supported by the (now) German Federal Ministry of Research, Technology and Space (BMFTR) under various funding schemes (ProteomeTools, 031L0008A; DIAS, 031L0168), the ERC grants (ORIGIN, 101077037) and (TOPAS, 833710), European Union’s Horizon 2020 Program (EPIC-XS, H2020-INFRAIA-2018-1, 823839), and the Elite Network of Bavaria (F-6-M5613.6.K-NW-2021-411/1/1).

## Author Contribution

M.W., D.P.Z, and B.K. conceived the study. W.G., D.P.Z., P.P., F.S., A.S., and G.M. designed experiments. D.P.Z, and F.P.B. performed laboratory experiments. W.G. trained different Prosit models. O.S. developed the integration with DLOmix. L.L. developed the integration with Koina. W.G. analyzed the data. W.G. and V.G. prepared visualizations. K.S., J.Z., T.K., U.R., and H.W. optimized, performed, and oversaw peptide synthesis. W.G. and M.W. wrote the manuscript. All authors reviewed, edited, and approved the manuscript.

## References

1. Beltrao, P., Bork, P., Krogan, N. J. & van Noort, V. Evolution and functional cross-talk of protein post-translational modifications. Mol. Syst. Biol. 9, 714 (2013).

2. Xu, H. et al. PTMD: A database of human disease-associated post-translational modifications. Genomics Proteomics Bioinformatics 16, 244–251 (2018).

3. Li, S., Iakoucheva, L. M., Mooney, S. D. & Radivojac, P. Loss of post-translational modification sites in disease. Pac. Symp. Biocomput. 337–347 (2010).

4. Han, Z.-J., Feng, Y.-H., Gu, B.-H., Li, Y.-M. & Chen, H. The post-translational modification, SUMOylation, and cancer (Review). Int. J. Oncol. 52, 1081–1094 (2018).

5. Meng, F., Forbes, A. J., Miller, L. M. & Kelleher, N. L. Detection and localization of protein modifications by high resolution tandem mass spectrometry. Mass Spectrom. Rev. 24, 126–134 (2005).

6. Doll, S. & Burlingame, A. L. Mass spectrometry-based detection and assignment of protein posttranslational modifications. ACS Chem. Biol. 10, 63–71 (2015).

7. Tyanova, S., Temu, T. & Cox, J. The MaxQuant computational platform for mass spectrometry-based shotgun proteomics. Nat. Protoc. 11, 2301–2319 (2016).

8. Kong, A. T., Leprevost, F. V., Avtonomov, D. M., Mellacheruvu, D. & Nesvizhskii, A. I. MSFragger: ultrafast and comprehensive peptide identification in mass spectrometry-based proteomics. Nat. Methods 14, 513–520 (2017).

9. Gessulat, S. et al. Prosit: proteome-wide prediction of peptide tandem mass spectra by deep learning. Nat. Methods 16, 509–518 (2019).

10. Wilhelm, M. et al. Deep learning boosts sensitivity of mass spectrometry-based immunopeptidomics. Nat. Commun. 12, 3346 (2021).

11. Gabriel, W. et al. Prosit-TMT: Deep learning boosts identification of TMT-labeled peptides. Anal. Chem. 94, 7181–7190 (2022).

12. Degroeve, S. & Martens, L. MS2PIP: a tool for MS/MS peak intensity prediction. Bioinformatics 29, 3199–3203 (2013).

13. Declercq, A. et al. Updated MS^2^PIP web server supports cutting-edge proteomics applications. Nucleic Acids Res. 51, W338–W342 (2023).

14. Zhou, X.-X. et al. PDeep: Predicting MS/MS spectra of peptides with deep learning. Anal. Chem. 89, 12690–12697 (2017).

15. Zeng, W.-F. et al. MS/MS spectrum prediction for modified peptides using pDeep2 trained by transfer learning. Anal. Chem. 91, 9724–9731 (2019).

16. Tarn, C. & Zeng, W.-F. PDeep3: Toward more accurate spectrum prediction with fast few-shot learning. Anal. Chem. 93, 5815–5822 (2021).

17. Bouwmeester, R., Gabriels, R., Hulstaert, N., Martens, L. & Degroeve, S. DeepLC can predict retention times for peptides that carry as-yet unseen modifications. Nat. Methods 18, 1363–1369 (2021).

18. Zeng, W.-F. et al. AlphaPeptDeep: a modular deep learning framework to predict peptide properties for proteomics. Nat. Commun. 13, 7238 (2022).

19. Picciani, M. et al. Oktoberfest: Open-source spectral library generation and rescoring pipeline based on Prosit. Proteomics 24, e2300112 (2024).

20. Zolg, D. P. et al. INFERYS rescoring: Boosting peptide identifications and scoring confidence of database search results. Rapid Commun. Mass Spectrom. 39 Suppl 1, e9128 (2025).

21. Buur, L. M. et al. MS2Rescore 3.0 is a modular, flexible, and user-friendly platform to boost peptide identifications, as showcased with MS Amanda 3.0. J. Proteome Res. 23, 3200–3207 (2024).

22. Zolg, D. P. et al. Building ProteomeTools based on a complete synthetic human proteome. Nat. Methods 14, 259–262 (2017).

23. Zolg, D. P. et al. ProteomeTools: Systematic characterization of 21 post-translational protein modifications by liquid chromatography tandem mass spectrometry (LC-MS/MS) using synthetic peptides. Mol. Cell. Proteomics 17, 1850–1863 (2018).

24. Gabriel, W. et al. Deep learning enhances precision of citrullination identification in human and plant tissue proteomes. Mol. Cell. Proteomics 24, 100924 (2025).

25. Creasy, D. M. & Cottrell, J. S. Unimod: Protein modifications for mass spectrometry. Proteomics 4, 1534–1536 (2004).

26. Simard, P. Y., Steinkraus, D. & Platt, J. C. Best practices for convolutional neural networks applied to visual document analysis. in Seventh International Conference on Document Analysis and Recognition, 2003. Proceedings 958–963 (IEEE Comput. Soc, 2005).

27. Kalyuzhnyy, A. et al. Profiling the human phosphoproteome to estimate the true extent of protein phosphorylation. J. Proteome Res. 21, 1510–1524 (2022).

28. The, M., MacCoss, M. J., Noble, W. S. & Käll, L. Fast and accurate protein false discovery rates on large-scale proteomics data sets with percolator 3.0. J. Am. Soc. Mass Spectrom. 27, 1719– 1727 (2016).

29. Fondrie, W. E. & Noble, W. S. Mokapot: Fast and flexible semisupervised learning for peptide detection. J. Proteome Res. 20, 1966–1971 (2021).

30. Ferries, S. et al. Evaluation of parameters for confident phosphorylation site localization using an Orbitrap Fusion tribrid mass spectrometer. J. Proteome Res. 16, 3448–3459 (2017).

31. Zong, Y. et al. DeepFLR facilitates false localization rate control in phosphoproteomics. Nat. Commun. 14, 2269 (2023).

32. Leijten, N. M., Heck, A. J. R. & Lemeer, S. Histidine phosphorylation in human cells; a needle or phantom in the haystack? Nat. Methods 19, 827–828 (2022).

33. Noberini, R. et al. Spatial epi-proteomics enabled by histone post-translational modification analysis from low-abundance clinical samples. Clin. Epigenetics 13, 145 (2021).

34. Wurtele, H. et al. Histone H3 lysine 56 acetylation and the response to DNA replication fork damage. Mol. Cell. Biol. 32, 154–172 (2012).

35. Jack, A. P. M. et al. H3K56me3 is a novel, conserved heterochromatic mark that largely but not completely overlaps with H3K9me3 in both regulation and localization. PLoS One 8, e51765 (2013).

36. Ding, N. et al. Chk1 inhibition hinders the restoration of H3.1K56 and H3.3K56 acetylation and reprograms gene transcription after DNA damage repair. Front. Oncol. 12, 862592 (2022).

37. Lange, U. C. et al. Dissecting the role of H3K64me3 in mouse pericentromeric heterochromatin. Nat. Commun. 4, 2233 (2013).

38. Di Cerbo, V. et al. Acetylation of histone H3 at lysine 64 regulates nucleosome dynamics and facilitates transcription. Elife 3, e01632 (2014).

39. Kacen, A. et al. Post-translational modifications reshape the antigenic landscape of the MHC I immunopeptidome in tumors. Nat. Biotechnol. 41, 239–251 (2023).

40. Sarkizova, S. et al. A large peptidome dataset improves HLA class I epitope prediction across most of the human population. Nat. Biotechnol. 38, 199–209 (2020).

41. Refsgaard, C. T., Barra, C., Peng, X., Ternette, N. & Nielsen, M. NetMHCphosPan - Pan-specific prediction of MHC class I antigen presentation of phosphorylated ligands. Immunoinformatics (Amst.) 1-2, 100005 (2021).

42. Solleder, M. et al. Deciphering the landscape of phosphorylated HLA-II ligands. iScience 25, 104215 (2022).

43. Will, A., Oliinyk, D., Bleiholder, C. & Meier, F. Peptide collision cross sections of 22 post-translational modifications. Anal. Bioanal. Chem. 415, 6633–6645 (2023).

44. Klaproth-Andrade, D., et al. Modanovo: A unified model for Post-translational modification-aware de Novo sequencing using experimental spectra from in vivo and synthetic peptides. *bioRxiv* 2025.09.12.675784 (2025) doi:10.1101/2025.09.12.675784.

45. Lautenbacher, L., et al. Koina: Democratizing machine learning for proteomics research. *bioRxivorg* (2024) doi:10.1101/2024.06.01.596953.

46. Zolg, D. P. et al. PROCAL: A set of 40 peptide standards for retention time indexing, column performance monitoring, and collision energy calibration. Proteomics 17, 1700263 (2017).

47. Shouman, O., Gabriel, W., Giurcoiu, V., Sternlicht, V. & Wilhelm, M. PROSPECT: Labeled tandem mass spectrometry dataset for machine learning in proteomics. Neural Inf Process Syst 35, 32882–32896 (2022).

48. Gabriel, W., Shouman, O., Schröder, E. A., Bößl, F. & Wilhelm, M. PROSPECT PTMs: Rich labeled tandem mass spectrometry dataset of modified peptides for machine learning in proteomics. Neural Inf Process Syst 37, 131154–131196 (2024).

49. Smith, L. N. Cyclical learning rates for training neural networks. in 2017 IEEE Winter Conference on Applications of Computer Vision (WACV) 464–472 (IEEE, 2017).

50. Lin, A., Plubell, D. L., Keich, U. & Noble, W. S. Accurately assigning peptides to spectra when only a subset of peptides are relevant. J. Proteome Res. 20, 4153–4164 (2021).

51. Fu, Y. & Qian, X. Transferred subgroup false discovery rate for rare post-translational modifications detected by mass spectrometry. Mol. Cell. Proteomics 13, 1359–1368 (2014).

52. Ramsbottom, K. A. et al. Method for independent estimation of the false localization rate for phosphoproteomics. J. Proteome Res. 21, 1603–1615 (2022).

53. Taus, T. et al. Universal and confident phosphorylation site localization using phosphoRS. J. Proteome Res. 10, 5354–5362 (2011).

54. Shteynberg, D. D. et al. PTMProphet: Fast and accurate mass modification localization for the Trans-Proteomic Pipeline. J. Proteome Res. 18, 4262–4272 (2019).

55. Tareen, A. & Kinney, J. B. Logomaker: beautiful sequence logos in Python. Bioinformatics 36, 2272–2274 (2020).

56. Sainburg, T., McInnes, L. & Gentner, T. Q. Parametric UMAP embeddings for representation and semisupervised learning. Neural Comput. 33, 2881–2907 (2021).

57. Malzer, C. & Baum, M. A hybrid approach to hierarchical density-based cluster selection. in 2020 IEEE International Conference on Multisensor Fusion and Integration for Intelligent Systems (MFI) 223–228 (IEEE, 2020).

