## Supplementary File for "Learning the Unseen: Data-Augmented Deep Learning for PTM Discovery with Prosit-PTM"

### Supplementary Figures

#### Supplementary Figure S1

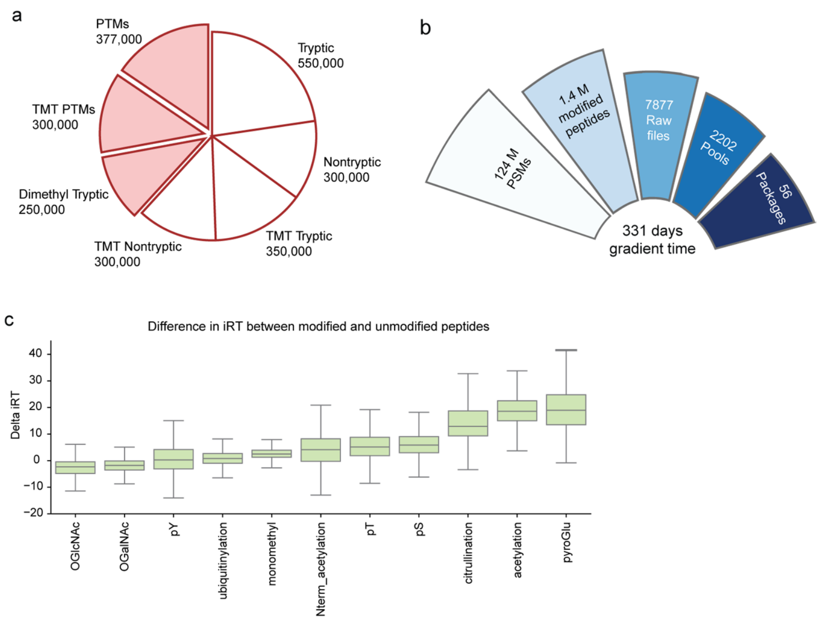

**ProteomeTools Dataset Composition.**

**a** Distribution of different peptide sets in Proteometools. The pie chart shows the distribution of synthesized modified peptides for each set.

**b** The dataset consists of 124 million PSMs derived from ~1.4M modified peptides extracted from 7877 raw files. Peptides were divided into 2202 across 56 different packages. The analysis required a total gradient time of 331 days.

**c** Box plots illustrate the differences in iRT between paired modified and unmodified peptides across eleven packages. The data show that modifications exhibit distinct, modification-specific chromatographic properties.

### Supplementary Figure S2

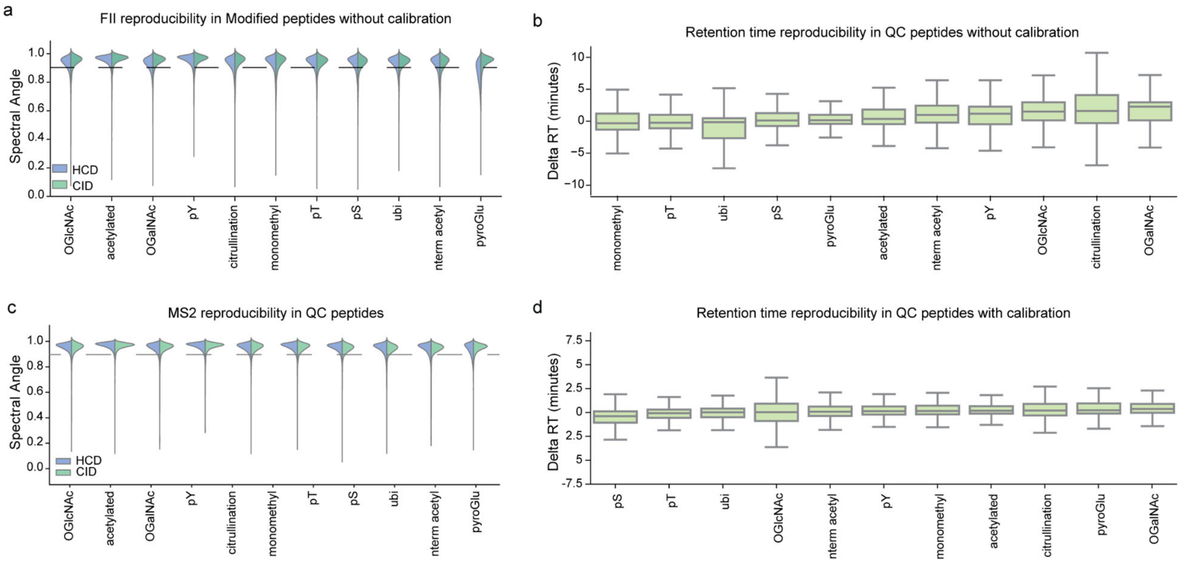

**Improved Spectral and Retention Time via Calibration.**

**a** Violin plots show the distribution of spectral angle (SA) values for quality control (QC) peptides without CE calibration, measured using HCD (orange) and CID (green) fragmentation methods. Different packages are represented along the x-axis.

**b** Box plots show the distribution of Retention Time values for quality control (QC) peptides without iRT calibration. Different packages are represented along the x-axis

**c** Violin plots show the distribution of spectral angle (SA) values for quality control (QC) peptides after CE calibration, measured using HCD (orange) and CID (green) fragmentation methods. Different packages are represented along the x-axis.

**d** Box plots show the distribution of Retention Time values for quality control (QC) peptides after iRT calibration. Different packages are represented along the x-axis.

### Supplementary Figure S3

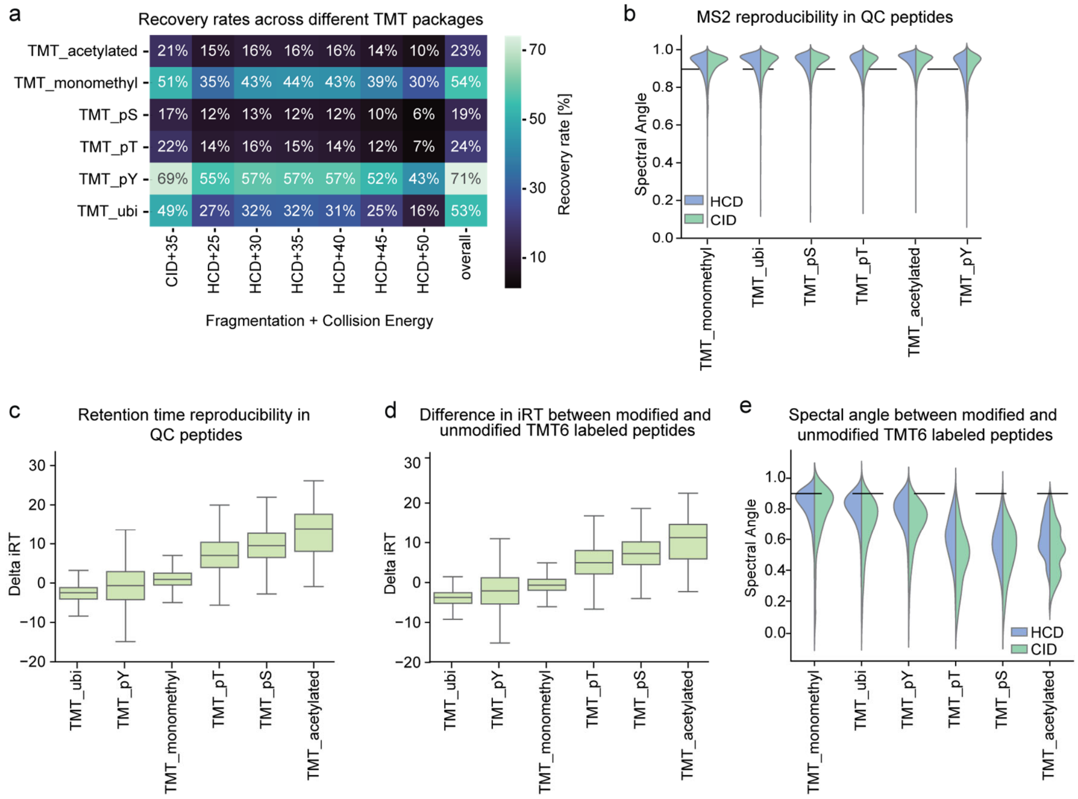

**Experimental Properties and Measurement Reproducibility of TMT-Labeled Modified Peptides.**

**a** The heatmap displays recovery rates (%) for different residue-ptm combinations (y-axis) for TMT6-labeled peptides analyzed under various mass spectrometric fragmentation conditions (x-axis).

**b** Violin plots show the distribution of spectral angle (SA) values for quality control (QC) peptides, measured using HCD (orange) and CID (green) fragmentation methods. Different packages are represented along the x-axis.

**c** Box plots show the distribution of Retention Time values for quality control (QC) peptides. Different packages are represented along the x-axis.

**d** Box plots illustrate the differences in iRT between paired modified and unmodified peptides across six packages. The data show that modifications exhibit distinct, modification-specific chromatographic properties.

**e** Violin plot compares FII between modified and unmodified peptides across different modifications and fragmentation methods (HCD, CID), highlighting the variability in the spectra, with some PTMs producing highly similar spectra and others leading to pronounced spectral divergence.

Supplementary Figure S4

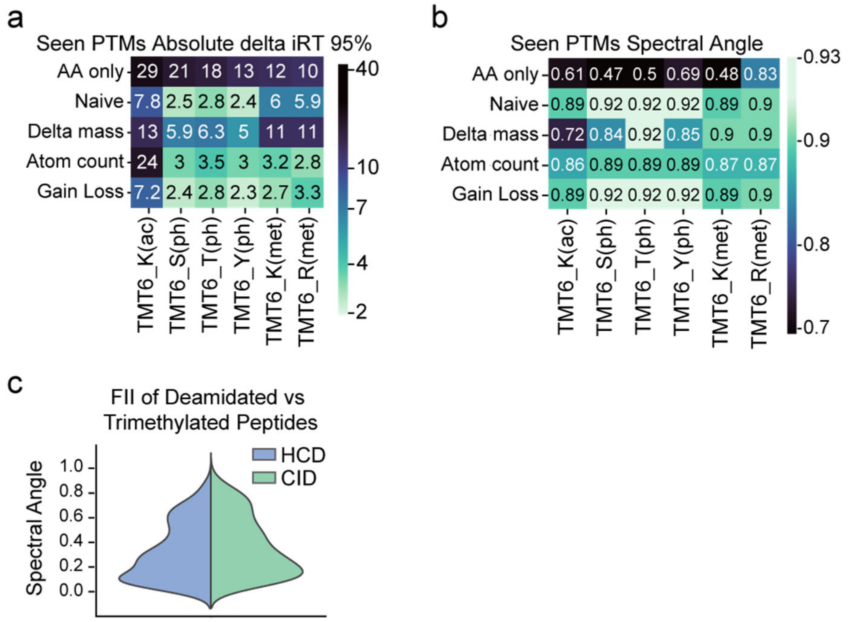

**Benchmarking PTM Prediction Performance on TMT-Labeled Peptides.**

**a** Comparative performance of PTM encoding techniques in Prosit for predicting modified TMT-6 labeled peptides retention times. Heatmap showing the 95th percentile delta iRT errors. Lower values (lighter green) indicate better predictions.

**b** Comparative performance of PTM encoding techniques in Prosit for predicting modified TMT-6 labeled peptides fragmentation patterns. Heatmap showing the Spectral angle between predicted and experimental spectra. Higher values (lighter green) indicate better spectral similarity.

**c** Violin plot compares FII between trimehtylated and acetylated peptides across different modifications and fragmentation methods (HCD, CID).

#### Supplementary Figure S5

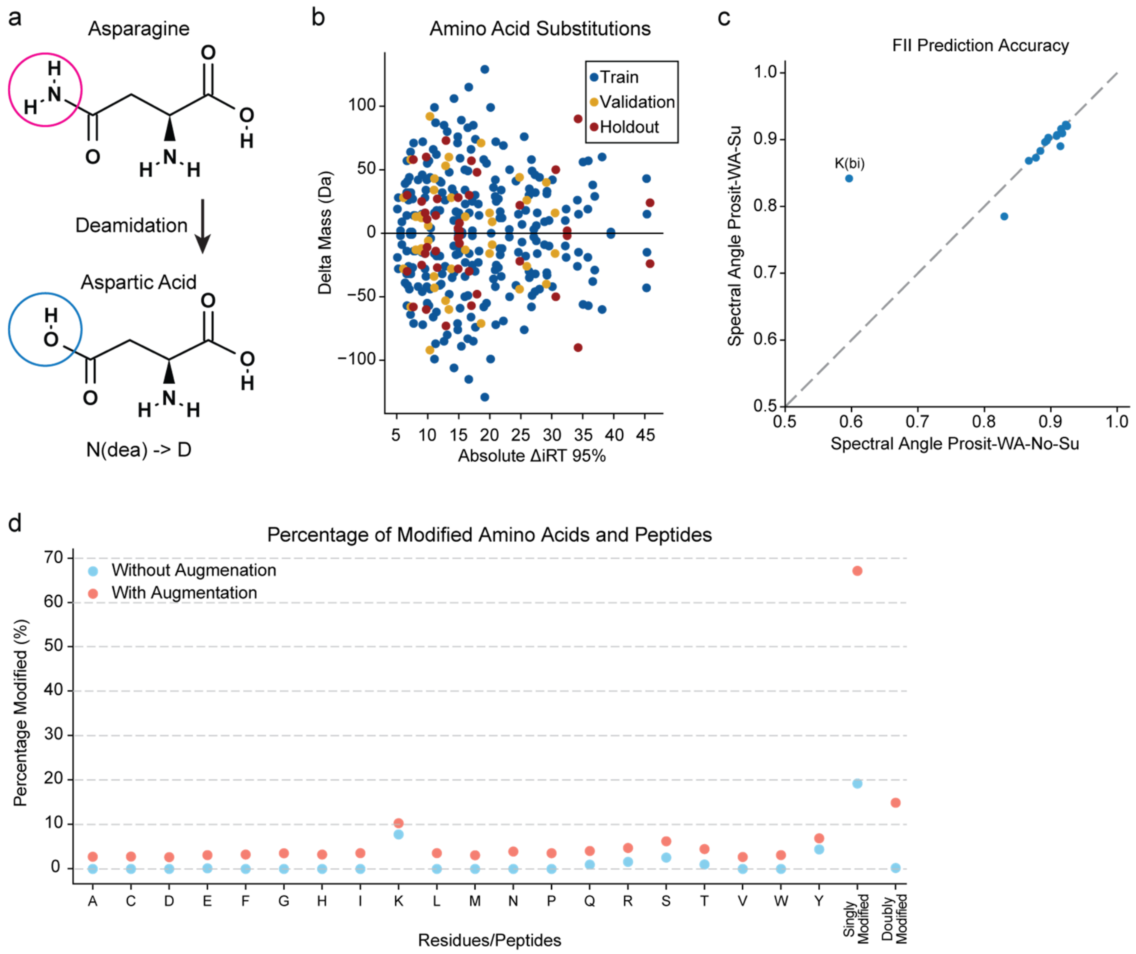

**Uniform Distribution of Modified Amino Acids Enabled by Augmentation Strategy.**

**a** The chemical structures illustrate the conversion of asparagine to aspartic acid via loss of an amide group, demonstrating a common post-translational modification that can lead to AAS.

**b** Percentage of modified AAs and peptides observed without augmentation (blue) and with augmentation using AAS (red). Each point represents the proportion of peptides with a given residue modified, as well as the overall percentage of singly and doubly modified peptides.

**c** Scatterplot shows the spectral angle for all unseen-PTMs with two prosit models: one trained on dataset with only AAS that contain sulfur atoms (Prosit-WA-Su) and another with all other AAS but the ones with sulfur atoms (Prosit-WA-No-Su). Lysine biotinylation is highlighted in the figure where Prosit-WA-Su outperforms Prosit-WA-No-Su.

**d** Scatter plot showing individual AAS, colored by dataset partition: green for training set, blue for validation set, and red for holdout set. The x-axis indicates the Delta iRT 95%, while the y-axis shows the delta mass (Da) associated with each substitution.

### Supplementary Figure S6

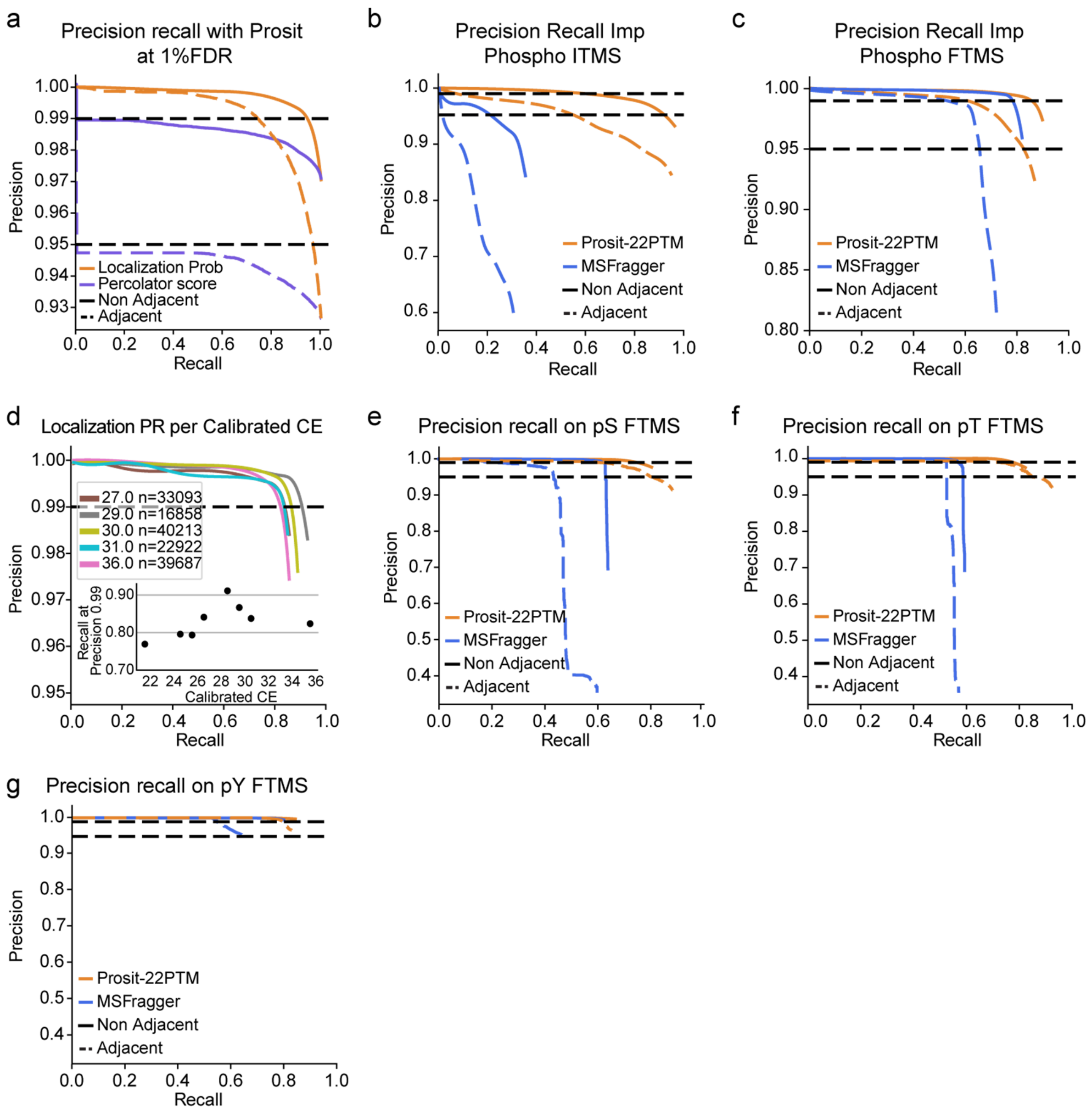

**Benchmarking Localization Probability for Phospho-Residue Assignment in Synthetic Data.**

**a** Precision-recall curve for phosphosite localization when using localization probability vs percolator score.

**b** Precision-recall curves for phosphosite localization of ITMS data for the important phosphosites package. Prosit-22PTM (orange) and MSFragger (blue) performance are shown, on nonadjacent (solid lines) and adjacent phospho residues (dashed lines). Black dashed lines indicate 95% and 99% precision thresholds.

**c** Precision-recall curves for phosphosite localization of FTMS data for the important phosphosites package. Prosit-22PTM (orange) and MSFragger (blue) performance are shown, on nonadjacent (solid lines) and adjacent phospho residues (dashed lines). Black dashed lines indicate 95% and 99% precision thresholds.

**d** Precision-recall curves demonstrating the impact of collision energy on phospho site localization performance for the important phosphosites package. Each curve corresponds to a different calibrated collision energy (CE) value ranging from 27.0 to 36.0. Each curve is labeled with its respective CE value and number of PSMs identified (n). With an inset scatter plot showing recall at 0.99 precision across different Calibrated CEs.

**e** Precision-recall curves for phosphosite localization of FTMS data for phosphosites on Serine only. Prosit-22PTM (orange) and MSFragger (blue) performance are shown, on nonadjacent (solid lines) and adjacent phospho residues (dashed lines). Black dashed lines indicate 95% and 99% precision thresholds.

**f** Precision-recall curves for phosphosite localization of FTMS data for phosphosites on Threonine only. Prosit-22PTM (orange) and MSFragger (blue) performance are shown, on nonadjacent (solid lines) and adjacent phospho residues (dashed lines). Black dashed lines indicate 95% and 99% precision thresholds.

**g** Precision-recall curves for phosphosite localization of FTMS data for phosphosites on Tyrosine only. Prosit-22PTM (orange) and MSFragger (blue) performance are shown, on nonadjacent (solid lines) and adjacent phospho residues (dashed lines). Black dashed lines indicate 95% and 99% precision thresholds.

#### Supplementary Figure S7

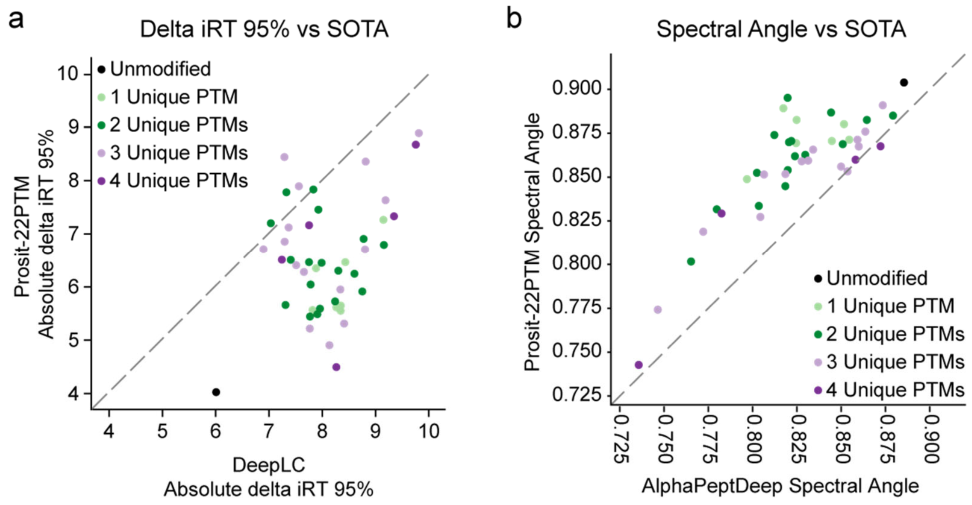

**Prosit-22PTM Outperforms State-of-the-Art for iRT and FII on Multiply Modified Peptides.**

**a** Scatterplot shows the Absolute delta iRT 95% for multiply modified peptides from ProteomeTools comparing Prosit-22PTM and DeepLC. Lower delta iRT 95% values indicate better prediction accuracy.

**b** Scatterplot shows the spectral angle (SA) for multiply modified peptides from ProteomeTools, comparing Prosit-22PTM and AlphaPeptDeep. Higher SA values indicate better prediction accuracy.

#### Supplementary Figure S8

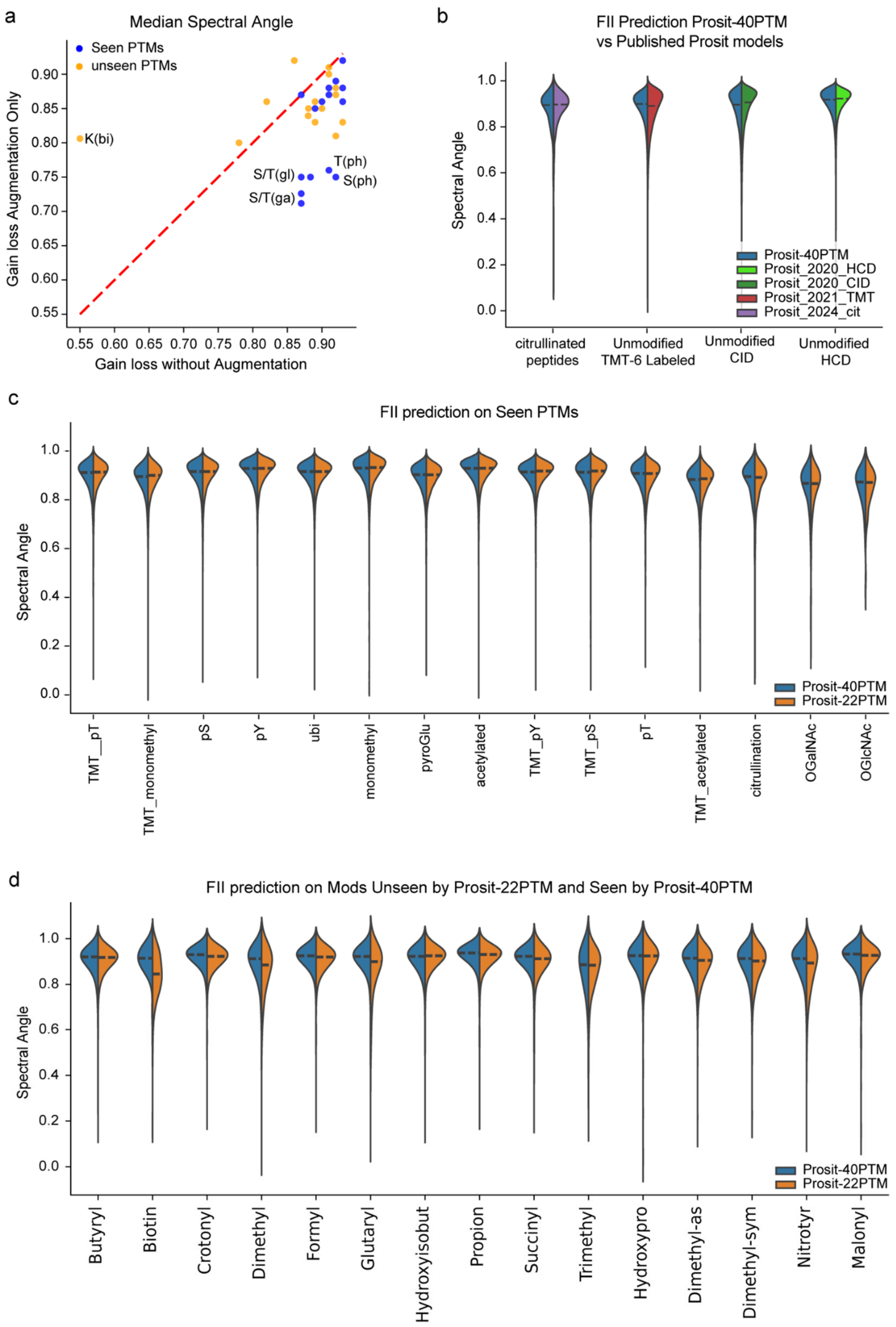

**Expanding PTM Coverage Using Only Amino Acid Substitution.**

**a** Scatterplot shows the spectral angle for all seen- and unseen-PTMs with Gain and Loss Prosit Model, and compares them to Prosit-2PTM. Blue dots indicate unseen-PTMs, and Yellow dots indicate seen PTMs.

**b** Violin plots show the distribution of spectral angle (SA) values comparing Prosit-40PTM vs published Prosit models. Different peptide types are represented along the x-axis.

**c** Violin plots show the distribution of spectral angle (SA) values comparing Prosit-40PTM vs Prosit-22PTM on seen PTMs. Different ProteomeTools packages are represented along the x-axis.

**d** Violin plots show the distribution of spectral angle (SA) values comparing Prosit-40PTM vs Prosit-22PTM on unseen PTMs by Prosit-22PTM and seen by Prosit-40PTM. Different ProteomeTools packages are represented along the x-axis.

### Supplementary Notes

#### Measurement Reproducibility Across ProteomeTools-PTMs

Previous studies have shown that acquiring the same sample multiple times can lead to technical variation, which may compromise reproducibility[1]. Additionally, the retention times (RT) of the same peptides can shift when injected with different samples due to matrix effects[2]. To monitor data consistency and reproducibility of ProteomeTools-PTM, PROCAL peptides[3] were spiked into each sample as calibration references across different acquisitions. We then evaluated the reproducibility of the fragment ion intensities (FIIs) across the Quality control (QC) peptides before and after calibration. Calibration of FII was applied only for HCD data, as CID collision energy is fixed and therefore not subject to calibration. For this, and all subsequent evaluations of FII, we use the normalized spectral contrast angle (short spectral angle, SA)[4], which evaluates the similarity between two vectors of FII. After calibration, we achieved an average SA of 0.94 for HCD fragmentation (Supplementary Figure S2c), representing a 0.04 improvement over the uncalibrated data (Supplementary Figure S2a). For CID fragmentation without calibration, the average SA was 0.93 across different PTMs. For indexed retention time (iRT), we observe that QC peptides elute within 0.5 minutes across different 60-minute gradient measurements after calibration (Supplementary Figure S2d), an improvement of 1.5 minutes compared to uncalibrated measurements (Supplementary Figure S2b).

#### Phospho localization Performance on Important Phospho Package

The important phospho package (TUM_imp_pSTY) comprised synthetic phosphopeptides with known biological functions, extracted from PhosphoSitePlus. This set was searched against the human proteome and acquired under similar conditions with multiple fragmentation settings. Prosit-22PTM again outperformed MSFragger, maintaining higher precision at the same recall level. First, we tested the pipelines on MS2 Spectra acquired with Iontrap; Prosit-22PTM achieved a recall of 0.8 at 99% precision, compared with a recall of 0.3 for MSFragger (Supplementary Figure S5b). When spectra were acquired with Orbitrap, Prosit-22PTM achieved a recall of 0.9 at 99% precision, compared with a recall of 0.82 for MSFragger (Supplementary Figure S5c).

Since ProteomeTools data were acquired using different NCE settings, we examined the effect of these settings on localization performance. We found that a calibrated NCE of 29 yielded the highest recall of 0.92 at 99% precision, consistent with previous phosphoproteomics analyses on tribrid mass spectrometers[5–7] (Supplementary Figure S6d). The resulting 29 collision energy is machine-dependent. We suggest users check the difference between their Mass Spectrometer's used collision energy and the calibrated one when predicting with Prosit-22PTM.

#### Comprehensive Training and Benchmarking of Prosit-40PTM Using Extended PTM Coverage

For completeness, we also trained Prosit-40PTM on all available 40 amino acid modifications in the updated ProteomeTools dataset, utilizing augmentation with the 342 AASs, which resulted in 382 amino acid-modification combinations in the training set. Prosit-40PTM performs on par with all earlier published Prosit models (Supplementary Figure S8b). Furthermore, for all PTMs in the updated ProteomeTools dataset, Prosit-40PTM achieves similar performance compared to Prosit-22PTM (Supplementary Figure S8c) and further increases prediction performance for biotinylation and trimethylation by ~0.05 SA each (Supplementary Figure S8d). Since Prosit-40PTM is either on par with or better than previous models, we believe Prosit-40PTM is ready for all use cases, as this generic model works very well on unmodified, modified, and labeled peptides. While its actual generalization performance cannot be tested at this time due to the lack of remaining synthetic peptides with unseen PTMs, we expect this model's generalization capability to be as good as for Prosit-22PTM because we used the same overall methodology for training (Methods).

#### Supplementary References

1. Piehowski, P. D. *et al.* Sources of technical variability in quantitative LC-MS proteomics: human brain tissue sample analysis. *J. Proteome Res.* **12**, 2128–2137 (2013).
2. Fang, N., Yu, S., Ronis, M. J. & Badger, T. M. Matrix effects break the LC behavior rule for analytes in LC-MS/MS analysis of biological samples. *Exp. Biol. Med. (Maywood)* **240**, 488–497 (2015).
3. Zolg, D. P. *et al.* PROCAL: A set of 40 peptide standards for retention time indexing, column performance monitoring, and collision energy calibration. *Proteomics* **17**, 1700263 (2017).
4. Toprak, U. H. *et al.* Conserved peptide fragmentation as a benchmarking tool for mass spectrometers and a discriminating feature for targeted proteomics. *Mol. Cell. Proteomics* **13**, 2056–2071 (2014).
5. Greguš, M., Koller, A., Ray, S. & Ivanov, A. R. Improved data acquisition settings on Q Exactive HF-X and fusion lumos tribrid Orbitrap-based mass spectrometers for proteomic analysis of limited samples. *J. Proteome Res.* **23**, 2230–2240 (2024).
6. Chang, A., Leutert, M., Rodriguez-Mias, R. A. & Villén, J. Automated enrichment of phosphotyrosine peptides for high-throughput proteomics. *J. Proteome Res.* **22**, 1868–1880 (2023).
7. Leutert, M., Rodríguez-Mias, R. A., Fukuda, N. K. & Villén, J. R2-P2 rapid-robotic phosphoproteomics enables multidimensional cell signaling studies. *Mol. Syst. Biol.* **15**, e9021 (2019).
